# *Syntrichia ruralis*: Emerging model moss genome reveals a conserved and previously unknown regulator of desiccation in flowering plants

**DOI:** 10.1101/2023.09.12.557352

**Authors:** Xiaodan Zhang, Jenna T. B. Ekwealor, Anderson T. Silva, Li’ang Yu, Andrea K. Jones, Brent D. Mishler, Andrew D. L. Nelson, Melvin J. Oliver

## Abstract

Water scarcity poses a significant threat to ecosystems in the face of global climate change. *Syntrichia ruralis*, a dryland moss known for its desiccation tolerance, provides valuable insights into surviving water-limited conditions. In this study, the genome of *S. ruralis* was sequenced and assembled into 12 chromosomes encompassing 21,169 protein-coding genes. Additionally, 3,199 unplaced scaffolds were identified as non-nuclear and symbiont DNA. Transposable elements (TEs) constitute 51.24% of the genome. Notably, chromosome 12, the largest in size due to its high TE load, was identified as the putative sex chromosome. Comparative analysis with the closely related *Syntrichia caninervis* genome reveals significant large-scale synteny yet some rearrangements, as well as the occurrence of older duplication events that are shared by both. Desiccation and drought tolerance associated gene families, such as early light-inducible proteins (ELIPs) and late embryogenesis abundant (LEA) proteins, were characterized. In addition to a subset of LEA genes being species-specific, a comparative transcriptomic analysis revealed that some shared LEA genes respond differently to dehydration in these two species. Many ELIPs (9 out of 30) are the product of tandem duplication events. As expected, our analyses revealed the importance of the phytohormone abscisic acid (ABA) in the desiccation response of *S. ruralis*. A significant number of ABA responsive genes were found to be regulated by *S. ruralis* orthologs of ABA insensitive 3 (ABI3) and abscisic acid responsive element binding factor 2 (AREB2). Markedly, an uncharacterized, but deeply conserved MYB transcription factor, appears to act as a negative regulator of AREB2 in *S. ruralis*. Interestingly, we determined that the orthologous MYB TF is also involved in an ABA-dependent stress response in the model flowering plant *A. thaliana*. In sum, the new genomic resources from this emerging model moss offer new insights into the evolution of desiccation tolerance in land plants.

## Introduction

Bryophytes (i.e. liverworts, mosses, and hornworts) hold a pivotal phylogenetic position as three of the four main living clades of land plants, along with the tracheophytes. As such, the study of bryophytes has been critical in establishing patterns that define the early evolution of plant development and growth (Naramoto et al., 2022). The origin of early land plants coincided with a large degree of genetic novelty (Bowles et al., 2020). A prime example includes the introduction of regulators (e.g., transcription factors) of critical phytohormone pathways such as abscisic acid (ABA), which itself has numerous roles governing drought and desiccation tolerance mechanisms. Since the emergence of land plants, these gene families have expanded, likely allowing for adaptation to increasingly diverse environments and developmental contexts (Donoghue et al., 2021). Indeed, desiccation tolerance mechanisms are believed to have been adapted to new developmental contexts as the need arose, such as late embryogenesis in seed plants (Oliver et al., 2005). Thus, understanding how desiccation tolerance is regulated in bryophytes can provide insights across land plant evolution.

Instrumental in understanding the significance of these genomic innovations has been the establishment of genetically tractable bryophyte models that exhibit desiccation tolerance. The tolerance of desiccation, which is the ability to equilibrate the water potential of tissues to that of the surrounding air (which is normally low) and then recover when water returns, is an important trait in land plant evolution. Bryophyte evolution spans the period when plants colonized the land from a freshwater origin (Mishler and Churchill, 1985); it has been postulated that vegetative desiccation tolerance was a critical requirement for plants to colonize terrestrial habitats (Oliver et al., 2005). However, the principal bryophyte model that is widely used to study growth and development, *Physcomitrium patens* (Lang et al., 2018; Rensing et al., 2020), is generally sensitive to dehydration stress and desiccation (Mishler and Oliver, 2009; Koster et al., 2010) and, although it has the capability for desiccation tolerance if treated with ABA (Oldenhof et al., 2006) or extremely slow drying rates in a laboratory setting (Greenwood and Stark, 2014; Xiao et al., 2018), it is not an ideal model for understanding tolerance to dehydration and desiccation.

Two closely related desiccation-tolerant mosses, *Syntrichia caninervis* and *Syntrichia ruralis* (previously known as *Tortula caninervis and Tortula ruralis*), have emerged as important model systems for understanding mechanisms of abiotic stress tolerance and their evolution. Both species have served as a focus for research into the ecological, reproductive, physiological, biochemical, and molecular aspects of desiccation tolerance in bryophytes (Proctor et al., 2007; Oliver 2009; Stark et al., 2005; Stark 2017; Coe et al., 2021; Zhang et al., 2011) and *S. caninervis* has proven amenable to genetic transformation (Li et al., 2023). *Syntrichia* is a diverse genus of mosses that occur in dryland habitats across the world and is one of the most ecologically dominant groups of mosses across western and northern North America (Brinda et al. 2021). Both *S. caninervis* and *S. ruralis* can dominate biological soil crust (biocrust) communities of North American drylands (Belnap and Lange 2003). However, whilst *S. caninervis* can be considered a dryland biocrust specialist, *S. ruralis* grows in a wider range of habitats that are not restricted to the biocrust community and span a range of moisture gradients (Mishler, 2007). Thus, despite their relatively recent divergence (∼ 5 MYA), the phenotypic variation within these two species argues that they are an excellent comparative resource for understanding how desiccation tolerance is experienced and regulated in land plants.

We report here the addition of a chromosomal-level genome assembly for *S. ruralis* and the insights this genome can provide to plant abiotic stress responses. Using comparative genomics and transcriptomics, we explored both evolutionary dynamics of the two *Syntrichia* genomes and the ways in which the two mosses regulate their molecular responses to desiccation and rehydration. We then expand our comparisons to flowering plants (i.e., *Arabidopsis thaliana*), where we used information from *S. ruralis* to identify a novel, but deeply conserved, ABA-associated transcription factor. Our results highlight the importance of developing bryophyte models and the conservation of desiccation tolerance associated pathways across land plants.

## Methods

### *S. ruralis* cultures

Shoots of *S. ruralis* (Hedwig), free of visible contamination with algae and bacteria, were vegetatively propagated on sterile fine sand collected from a dune near Moab, Utah (93.9 % sand, 5.5% silt, 0.6% clay with a of pH 8.4), and watered on alternating weeks with sterile distilled water and with a 30% inorganic nutrient solution (Hoagland & Arnon 1938). The shoots originated from a single specimen (Brinda 9108) collected from the Bow River Valley in Calgary Alberta (51.098875, −114.281461) and vouchered in the University of Nevada, Las Vegas herbarium. Cultures were placed in a growth chamber set to a 12-hr photoperiod (20°C light, 8°C dark), at ∼90 µmol m^-2^ s^-1^ photosynthetically active radiation (PAR). The single clonal line used for genome sequencing had been sub-cultured and grown to maturity through at least five asexual generations. Several gametophytes from the original clonal line were used to expand and establish subcultures for generating sufficient material for isolation of RNA for construction of transcriptome libraries for transcript abundance studies (RNAseq). Expansion of cultures was achieved by isolating individual shoots following branching as well as by fragment regeneration.

Desiccation (slow drying) was achieved by placing the gametophytes in small wire baskets over saturated ammonium nitrate (67% relative humidity) in a closed glass desiccator placed in the same incubator as the moss cultures at 20°C. Under these conditions the gametophytes reach a stable weight (equilibrium) at 6 h and a water potential of −54 MPa within the light period (100 μmol m 2 s 1) of the day/night cycle. Rehydration was achieved by placing the desiccated gametophytes in a petri dish in the incubator at 20°C in the light and adding sufficient distilled water to ensure full hydration.

### Genomic DNA isolation and Chicago library preparation and sequencing

Genomic DNA isolation, library preparation, sequencing, and assembly were conducted by Dovetail Genomics (Scotts Valley, CA) and as described for the *S. caninervis* genome (Silva et al, 2020). Briefly, genomic DNA was isolated from 1-2 grams of powdered flash frozen gametophytes using a standard CTAB-based procedure (Silva et al., 2020) and high molecular weight genomic DNA was precipitated, resuspended in Qiagen Buffer G2 with RNAse A and after incubation at 50°C for 30 minutes, the DNA was purified using Qiagen Genomic-tips (Qiagen Waltham, MA). Chicago genomic DNA libraries were prepared as described by Putnam et al, 2016. The libraries were sequenced on an Illumina HiSeq platform to produce 389 million 150 bp paired-end reads, which provided 153.2 x physical coverage of the genome (1-100 kb pairs). A Dovetail HiC library was prepared as described by Erez Lieberman-Aiden et al., 2009. For each library, chromatin was fixed in place in the nucleus by incubation of the gametophytes in 1% formaldehyde for 15 minutes under vacuum. The fixed chromatin was extracted from the treated gametophytes using the DovetailTM Hi-C Kit (Dovetail Genomics, Santa Cruz, CA). After digestion with DpnII, the 5’ overhangs filled in with biotinylated nucleotides and ligated. After ligation, crosslinks were reversed, and the DNA purified from protein. The DNA was then sheared to ∼350 bp mean fragment size and sequencing libraries were generated using NEBNext Ultra enzymes and Illumina-compatible adapters. Biotin-containing fragments were isolated using streptavidin beads before PCR enrichment of each library. The libraries were sequenced on an Illumina HiSeq platform to produce 224 million 2×150bp paired end reads, which provided 23,246.36 x physical coverage of the genome (10-10,000 kb pairs).

A de novo assembly was constructed using a combination of paired-end (mean insert size ∼350 bp) libraries. De novo assembly was performed using `Meraculous` v2.2.2.5 (diploid mode 1) (Chapman et al, 2011) with a k-mer size of 109. The input data consisted of 385.9 million read pairs sequenced from paired end libraries (totaling 112.9 Gbp). Reads were trimmed for quality using Trimmomatic (Bolger et al 2015). The input de novo assembly, shotgun reads, Chicago library reads, and Dovetail HiC library reads were used as input data for HiRise, a software pipeline designed specifically for using proximity ligation data to scaffold genome assemblies (Putnam et al, 2016) as described by Silva et al., 2020.

### Repeat annotation

`RepeatModeler` (v1.0.11) (http://www.repeatmasker.org/RepeatModeler) and `LTR_retriever` (Ou and Jiang 2018) (which was employed to identify intact LTRs) pipelines were used to de novo identify repeat element families in the *S. ruralis* genome assembly. The de novo repeats that matched known *P. patens* or *S. caninervis* genes (`BLASTN` E-value < 1 × 10^-5^) were removed from the LTR library. The remaining unclassified sequences were subsequently curated using a `BLASTN` comparison with a characterized plant protein database (alluniRefprexp082813). The resultant de novo library was combined with the RepBase plant repeat database (Bao et al., 2015) for input into `RepeatMasker` (v4.0.7) to discover and identify repeats.

### Genome annotation

The *S. ruralis* genome was annotated using `MAKER` (2.31.8) (Canterel et al., 2008) and the *ab initio* was performed using `SNAP` (Zaharia et al., 2015), Augustus (Stanke and Morgenstern 2005), and `BRAKER` (v2.1.2. Hoff et al., 2019) with two and one round of reiterative training, respectively. The de novo transcriptome assembly was treated as expressed sequence tag evidence and protein sequences from *Arabidopsis thaliana*, *Oryza sativa*, *P. patens*, *S, caninervis* and UniprotKB plant databases were used as protein homology evidence for `MAKER`. The custom LTR library was used as input as a custom repeat database in addition to the default protein repeat libraries in `MAKER`. The initial iteration of the `MAKER` gene set was filtered for gene models with an Annotation Editing Distance (AED) score equal to 0.25 and only the gene models with more than 50 aa were retained.

The 3199 scaffolds that were not assembled into chromosomes were considered as symbiont-related DNA sequences because they were alike to one another based on the link-density map and had much higher read coverage, as expected for contaminants. To validate our prediction that the unplaced scaffolds were symbiont-derived, the 5,896 associated genes were used as queries in a BLAST against the NCBI non-redundant nucleotide database (O’Leary et al., 2016).

### Syntenic comparisons between genomes of *S.ruralis*, *S. caninervis* and *C.purpureus*

To identify syntenic gene blocks between *S. ruralis*, *S. caninervis,* and *C. purpureus*, an all-against-all `BLASTP` analysis was conducted, using a stringent threshold (E value < 1 × 10^-5^ and top five matches). Syntenic blocks were defined based on the presence of a minimum of five synteny gene pairs, employing the `MCScanX` package with default settings (Wang et al., 2012). Subsequently, adjacent blocks were merged, and large syntenic blocks were chosen for further analysis. These sizable syntenic blocks were utilized to infer the chromosome evolution and explore the syntenic relationships between *S.ruralis*, *S. caninervis* and *P. patens*.

### Identification and comparison of gene families among different species

Protein sequences of 11 other plant species were downloaded from the NCBI database (https://www.ncbi.nlm.nih.gov/) with the exception of protein sequences for *O. thomaeum* that were kindly provided by Dr. Robert Vanburen, Michigan State University. The species include (1) three species with the vegetative DT at most environmental conditions: *S. ruralis*, *S. caninervis*, and *O. thomaeum*; (2) two species with vegetative DT only at certain environmental conditions: *M. polymorpha* and *P. patens* (Koster et al., 2010; Greenwood et al., 2014); and (3) seven species without vegetative DT but with desiccation-tolerant seeds or spores: *A. filiculoides*, *A. thaliana*, *A. trichopoday*, *C. quinoa*, *O. sativa*, *S. cucullata* and *S. moellendorffii*. Identification and comparison of gene families were conducted using `OrthoFinder2` using default parameters (Emms and Kelly, 2019).

### Identification and phylogenetic analysis of LEA and ELIP genes

LEA and ELIP genes were obtained from *S. ruralis* and *S. caninervis* by `HMMER` (http://hmmer.org/) and NCBI `BLAST`. HMM profiles of eight LEA subfamilies (LEA_1, PF03760; LEA_2, PF03168; LEA_3, PF03242; LEA_4, PF02987; LEA_5, PF00477; LEA_6, PF10714; DHN, PF00257 and SMP, PF04927) were retrieved from the Pfam database (http://pfam.xfam.org) and used to identify LEA domain-conserved proteins using the hmmsearch program. HMM profiles of ELIPs were downloaded from the interpro database (https://www.ebi.ac.uk/interpro/). Proteins identified with hmmsearch were BLAST against the nr database to confirm their LEA-domain classifications.

The amino acid sequences of 59 *S. ruralis* and 56 *S. caninervis* LEAs were aligned using `Clustal Omega`. Then the phylogenetic tree was constructed by Molecular Evolutionary Genetic Analysis software (Mega5), the Maximum Likelihood (ML) method, and the tree was visualized by Interactive Tree of Life online tool (iTOL, https://itol.embl.de/itol.cgi).

### RNA extraction and RNA sequencing

RNA was extracted using the RNeasy (Qiagen, Hilden, Germany) kit with the RLC buffer following the manufacturer’s recommended protocol. RNA isolates were treated with DNase1 and cleaned using the DNA-free RNA Kit (Zymo Technologies, Irvine, CA). RNA quality was assessed by use of a fragment analyser (Advanced Analytical Technologies, Ankeny, IA) and concentration determined by use of a Nanodrop Spectrophotometer (ThermoFisher, Waltham, Massachusetts). RNA libraries were created and individually barcoded from 2.7 µg of template total RNA utilizing the TruSeq RNA Sample Prep Kit (Illumina, San Diego, CA) as described in the manufacturer’s recommended protocol. Libraries were pooled in groups of twelve and sequenced (twelve samples per lane) on an Illumina HiSeq 2500 ultra high-throughput DNA sequencing platform (Illumina, San Diego, CA) at the DNA Core facility at the University of Missouri, Columbia, MO, USA (http://dnacore.missouri.edu//HiSeq.html).

### Transcriptome analysis

The RNA-seq data was processed using the analysis suite `RMTA` (Read Mapping Transcript Assembly), which involved trimming the adapter and eliminating low-quality sequences from the raw data, as well as mapping the trimmed sequences to the genome and quantifying transcript levels in read counts (Peri et al, 2020). Differential abundance of transcripts was tested by the `DESeq2` package (Anders, 2010) in `R` (Love, Michael et al, 2014). Transcripts with adjusted *P*-valuex < 0.05 and |log2(fold change)| > 1 were considered to be differentially abundant. Venn diagrams of genes with differential transcript abundance between treatments were displayed by the `R` package `ggVennDiagrams` (Gao, Chun-Hui, 2021). Furthermore, gene ontology (GO) and enrichment analysis were performed to explore the functional characteristics of genes exhibiting differential RNA abundance using TopGO (Alexa, A., & Rahnenführer, J. 2009).

### Identification of transcription factors regulating the desiccation response in *S. ruralis*

Transcription factors in *S. ruralis* were identified through `PlantTFDB` (Tian et al., 2020). The 2 Kb upstream region of the 3,045 genes found to be differentially abundant between control and SD conditions was extracted and used as the input data for performing AME (Analysis of Motif Enrichment; McLeay RC et al., 2010) with TF-DNA binding data from the Arabidopsis DAP motif database (O’Malley et al., 2016) to identify overrepresented motifs and corresponding *S. ruralis* TFs. Pairwise abundance correlation was calculated between these corresponding *S. ruralis* TFs and putative target genes across all stress-associated conditions. Regulatory network of TFs and putative targets was generated by Cytoscape (Saito, Rintaro et a.l, 2013)

### Genotyping of *A. thaliana* T-DNA insertion line and ABA treatment

Genomic DNA was extracted from young leaves of T-DNA *Atmyb55* mutant (CS823517) using the CTAB-based method as described by Doyle and Doyle (1987) with minor modifications. Primers were designed using T-DNA primer design website (http://signal.salk.edu/tdnaprimers.2.html) and used to PCR amplify and genotype the mutant. Wild type (Col-0) and *Atmyb55* homozygous mutants on MS-agar plates with concentrations of ABA (0, 0.1, 0.3 and 0.5 uM) and monitored germination every 12 hours for six days. Wild type and *Atmyb55* seedlings grown on 0 uM ABA for one week were transferred to plates containing 2 uM ABA and monitored for variation in root growth for an additional three days.

## Results & Discussion

### De novo assembly of *Syntrichia ruralis* genome

The draft genome of *S. ruralis* was assembled into 381.24 Mb, which consisted of 3,211 scaffolds with an N50 value of 24.06 Mb. The twelve largest scaffolds, representing 80% of the assembly (305.19 out of 381.24 Mb), were considered near-chromosome level assemblies, with the consensus plant telomere repeat (TTTAGGG, Fajkus et al., 2019; Fajkus et al., 2021) present at 8 of the 24 scaffold ends. The additional 3,199 unplaced scaffolds were determined to be putative symbiont-derived or plastid gDNA (Table 1). The size of the 12 *S. ruralis* chromosomes ranged from 15.7 to 48.2 Mb (**Table S1**), with the largest being the putative sex chromosome (Chr. 12; **Figure 1**).

**Figure 1.**
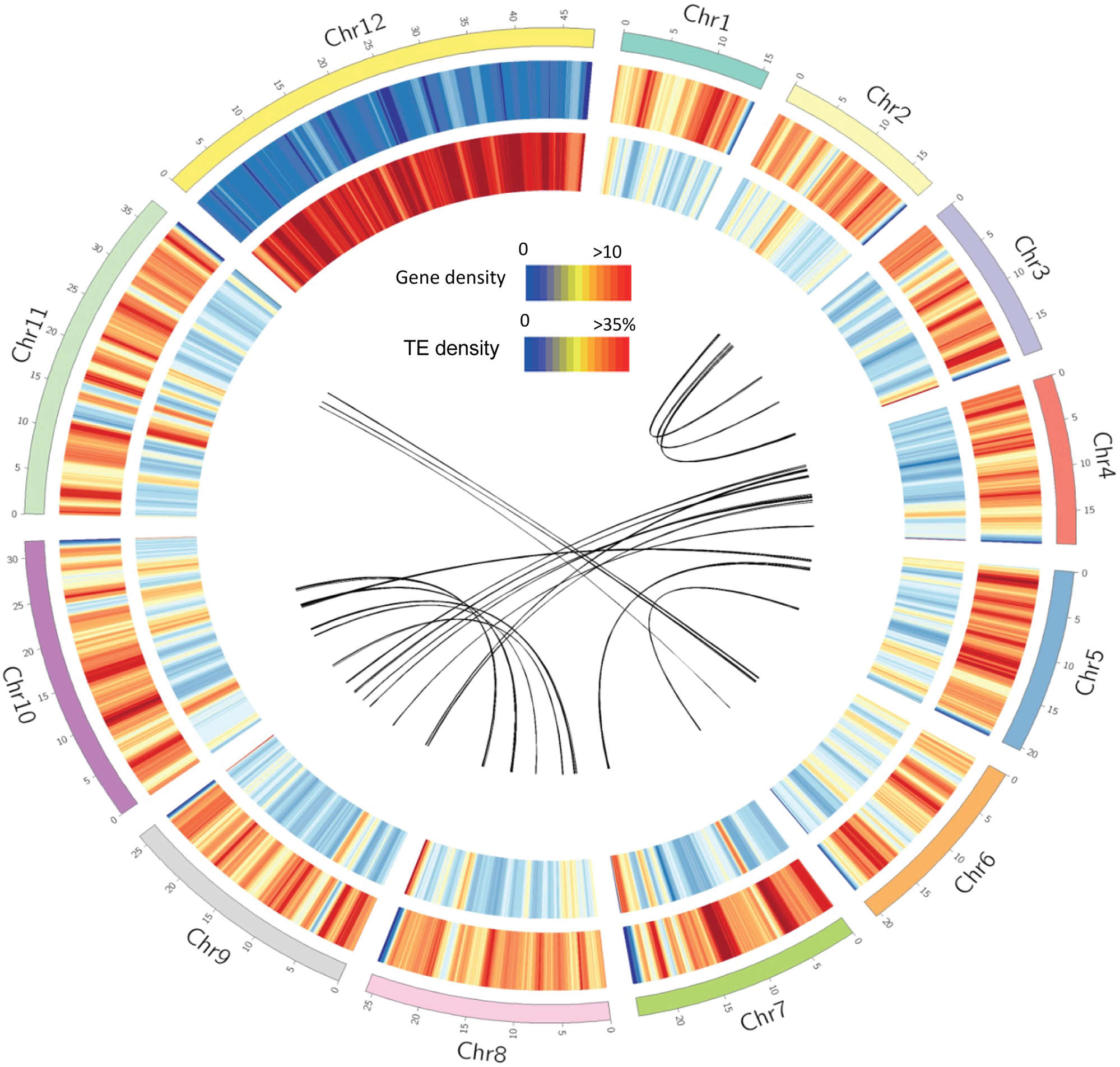
Characteristics of the *Syntrichia ruralis* genome. From outer to inner: The 12 chromosomes; gene density in 100-kb genomic regions; transpose element (TE) density in 100-kb genomic regions; inter-chromosome genomic collinearity.

The *S. ruralis* reference genome represents the fourteenth publicly-available moss genome, and so far all have small genomes, relative to vascular plants (**Figure S1**). The estimated genome size, 305.19 Mb, with a nuclear GC content of 44.6%, was smaller than that of closely-related *S. caninervis* (331.83 Mb) and consists of 12 rather than 13 chromosomes (Silva et al., 2020) and is also smaller than that of the distantly-related *Takakia lepidozioides* (331 Mbp; Hu et al., 2023). Similarly, the *S. ruralis* genome is smaller than that of the first, and most highly characterized model moss, *Physcomitrium patens* (475.8 Mb; Lang et al., 2018; Rensing et al., 2020). The genome size of *Ceratodon purpureus*, a moss in the same taxonomic subclass (Dicranidae), also has a larger genome (358 Mb male genome with 13 chromosomes; Carey et al., 2021), as do three more distantly-related pleurocarpous mosses with recent genome assemblies: *Entodon seductrix* (348.4 Mb; Yu et al., 2022), *Fontinalis antipyretica* (385.2 Mb; Yu et al., 2020), *Calohypnum plumiforme* (434 Mb; Mao et al., 2020), and the very large Bryidae moss *Pohlia nutans* (698.20 Mbp; Liu et al., 2022). However, the genome of a fourth pleurocarpous moss, *Hypnum curvifolium*, is estimated to be smaller than that of *S. ruralis,* at 262.0 Mb (Yu et al., 2022). The 305.19 Mb *S. ruralis* genome is similar in size to that of *Bryum argenteum* (292.04 Mb; Gao et al., 2023) and *Pleurozium schreberi* (318 Mb; Pederson et al., 2019). Of the two publicly-available *Sphagnum* genomes, one assembly is similar in size to *S. ruralis*, *Sphagnum angustifolium* at 367 Mb, while the other, *Sphagnum divinum*, is larger at 424 Mb (Healey et al., 2023). In sum, the size of the *S. ruralis* genome is within the range of other mosses, and there does not appear to be much of a pattern with respect to genome size in moss taxonomic groups as each are variable.

A total of 27,065 genes were predicted in the draft genome including 21,169 gene models associated with the 12 chromosomes and 5,896 genes with the 3,199 symbiont-derived scaffolds (**Table 1**). The number of genes per chromosome ranged from 857 to 2,888. Despite being the largest, Chr. 12 had the fewest predicted genes (**Table S1**, **Figure 1**). To validate our prediction that the unplaced scaffolds were symbiont-derived, the 5,896 associated genes were used as queries in a BLAST against the NCBI non-redundant nucleotide database (O’Leary et al., 2016). Significant hits were recovered for 5,765 (97.8%) genes. Of these, 5,272 genes (91.45%) were bacterial-associated, 343 genes (5.95%) were plant-related, and the rest (150, or 2.5%) were associated with fungi and single-cell eukaryotes (**Table S2**). Among the 21,169 protein-coding genes in *S. ruralis* chromosomes, 5,172 genes were observed in a tandem repeat configuration of 2 to 25 genes (**Table S3**). Chromosome 11 exhibited the most tandemly arrayed gene clusters, while no tandemly duplicated gene clusters were observed on Chr. 12.

Though the *S. ruralis* genome is ∼ 8% smaller than that of *S. caninervis, S. ruralis* has almost 30% more genes: 21,169 in *S. ruralis* vs. 16,545 in *S. caninervis*, **(Figure S1**). (Silva et al., 2020). Among published moss genomes, the total number of predicted genes in *S. ruralis* is intermediate between *S. caninervis* (16,545), *P. schreberi* (15,992), *F. antipyretica* (16,538), and *B. argenteum* (17,721) on the low end, and *H. curvifolium* (29,077), *Physcomitrium patens* (32,926), and *Pohlia nutans* (40,905) on the high end (SIlva et al., 2020; Pederson et al., 2019; Yu et al., 2020; Gao et al., 2023; Yu et al., 2022; Lang et al., 2018; Rensing et al., 2020; Liu et al., 2022).

Transposable elements identified by the `EDTA` (Extensive De novo Transposable element Annotator; Ou et al., 2019) pipeline revealed the presence of diverse transposable elements (**Table S4, Figure S2**). The identified TEs (51.24%) were categorized into different classes, including class I retrotransposons (8.12%), class II DNA transposons (35.42%), and unclassified repeat regions (7.58%). Class I retrotransposons were sub grouped into non-LTR retrotransposons (0.13%) and LTR retrotransposons (7.99%). Class II DNA transposons included TIR transposons (19.69%), nonTIR (15.73%), and unclassified DNA transposons (0.11%). Interestingly, unlike most plant genomes where LTR retrotransposons are the most abundant types, class II DNA transposons (TIR) were the most abundant in *S. ruralis* genome. LTR-Gypsy (47.9 %) was the most abundant type in *P. patens* genome (Lang et al., 2018; Rensing et al., 2020), while LTR-Gypsy only accounted for 3.59% in *S. ruralis* genome. Unknown repeat element was the most abundant type in the *S. caninervis* genome, representing 36.8%, in comparison to 7.58% in the *S. ruralis* genome (Silva et al., 2020). The differences in TE structures among these three genomes could be associated with the difference in response to the desiccation tolerance. The TEs were not evenly distributed across the *S. ruralis* genome, ranging from 46.3% to 64.7%, with the Chr. 12 (the sex chromosome; see below) harboring the highest TE percent (**Table S5, Figure 1**). The high percentage of masked regions suggests that TEs play a significant role in shaping the genome structure and dynamics.

### Evolution of the *S. ruralis* genome

Conservation of genome structure and collinearity between species can provide insight into major evolutionary events over history. First, CoGe’s `SynMap` tool (Lyons, E. and Freeling, M., 2008) was used to assess the degree of genome conservation between close relatives *S. ruralis* and *S. caninervis*, which last shared a common ancestor ∼5 MYA (Jauregui-Lazo et al., 2023; **Figure 2A**). In addition to some smaller inversion events along *S. ruralis* Chrs. 1, 2, 9, and 10, this pairwise comparison also uncovered a chromosomal fusion or fission event between *S. ruralis* Chr. 11 and *S. caninervis* chromosomes 2 and 8 (**Figure 2A**). A closer examination of the genes falling within these inversion events reveals a number of heat shock and stress-responsive transcription factors with different responses to desiccation tolerance between *S. ruralis* and *S. caninervis* (**Table S6**). To infer the directionality in this chromosomal rearrangement, both mosses *Ceratadon purpureus* and *Physcomitrella patens* were added to 3-way genome-wide synteny comparisons with `McScan` (**Figure 2B and Figure S3;** *P. patens*). The inclusion of *C. purpureus* as an outgroup, which is reported to have last shared a common ancestor with *Syntrichia* ∼180 MYA (Jauregui-Lazo et al., 2023) and thus is more closely related to *S. ruralis* than is *P. patens* (Zhong et al., 2014), revealed that Chr. 11 in *S. ruralis* likely resulted from a chromosomal fusion event of *S. caninervis* Chrs. 2 and 8 sometime after these two species last shared a common ancestor. These data highlight the different genomic landscapes that have emerged in these two desiccation tolerant mosses.

**Figure 2.**
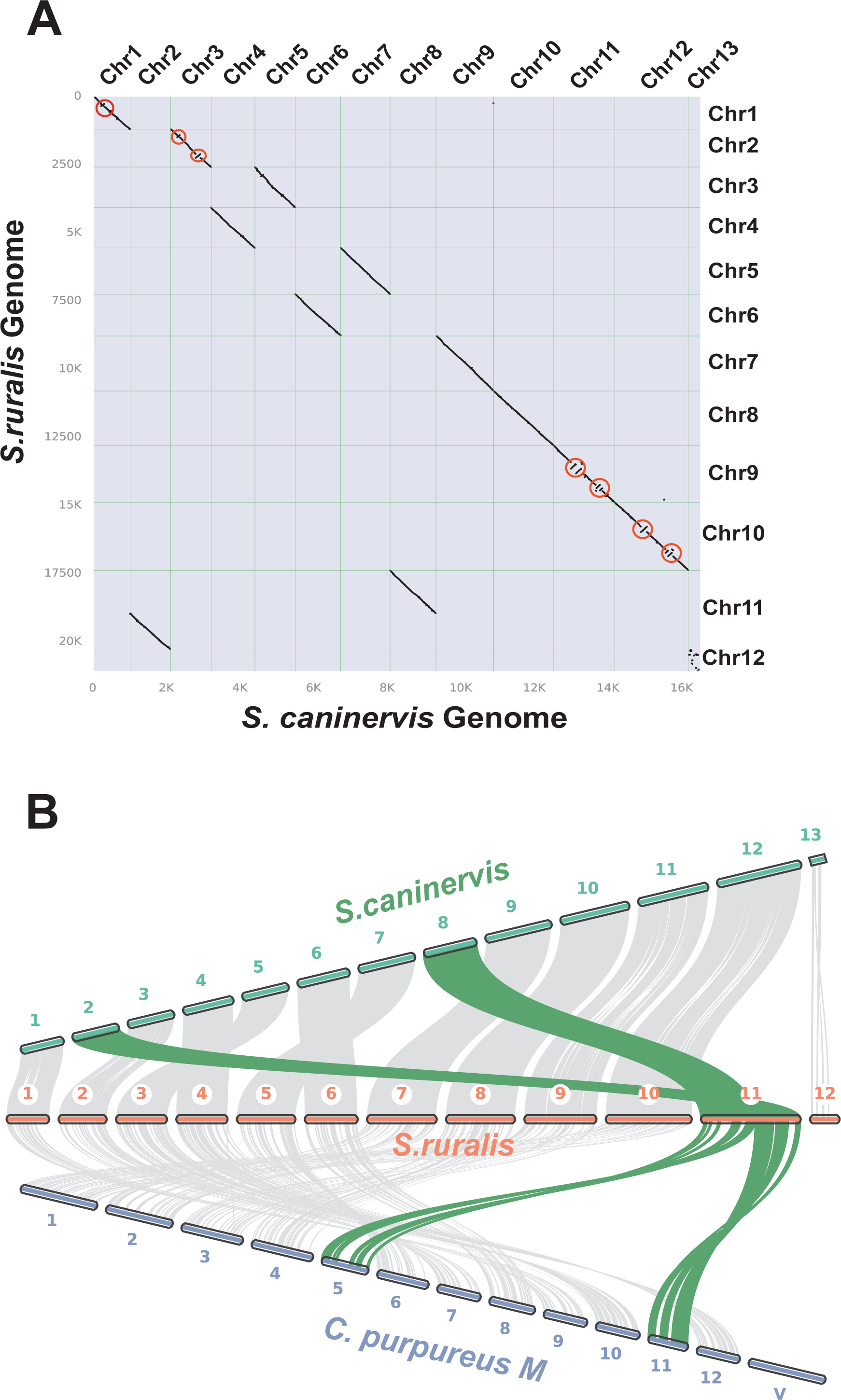
Comparative genomics between *S. ruralis, S.caninervis* and *C.purpureus*. **A.** Dot plot graph of genome synteny revealed genome duplication and small-scale inversions between *S. ruralis* and *S. caninervis*. Several inversions (highlighted in red circles) were observed in *S. ruralis* chromosomes 1, 2, 9 and 10. **B.** Large-scale genome conservation between *S. ruralis*, *S. caninervis* and *C.purpureus* generated by McScan. Each node (connection point) represents a genomic region, and the edges (lines) connecting the nodes represent syntenic blocks or collinear regions of syntenic genes. The length and thickness of the edges indicate the size and similarity of the syntenic blocks, respectively. Conserved syntenic blocks are depicted in shared colors. The green edges reflect a chromosomal rearrangement event (fusion event) between *S. caninervis* Chr. 2 and 8 and *S. ruralis* Chr. 11.

In *S. ruralis*, Chr. 12, which is the largest in size (48.2 Mb; ∼1.3 times bigger than the largest autosome) but has the fewest number of genes (857 vs. 2,888 in Chr. 11), is proposed to be the putative sex chromosome, specifically the V (male) chromosome. Sex-specific motifs from *S. caninervis* (gene id: Sc_g00229) were mapped to gene Sr_g00673 on Chr. 12 with 88% identity (e-value = 0), supporting this conclusion. Comparative analysis revealed that there were significantly fewer syntenic blocks between the U and V sex chromosomes compared to the autosomes of both *S. ruralis* and *S. caninervis*, suggesting the presence of sex-specific genes. While the observed lack of synteny may be due to algorithmic difficulty of `McScan` in aligning the repetitive sequence of these two chromosomes, a separate examination of the protein coding sequences using an all-versus-all protein `BLAST` (`BLASTp`) (e-value = 10e-10, identity > 80%) revealed a small portion of homologous genes between the sex chromosomes of *S. ruralis* and *S. caninervis* (139 out of 857 genes in *S. ruralis*) (**Table S7**), highlighting the stark differences between these two chromosomes. In *C. purpureus,* the U and V sex chromosomes share no obviously syntenic regions with each other or with other *C. purpureus* autosomes (Carey et al., 2021). This suggests that the male V sex chromosome of *S. ruralis* and the female U chromosome of *S. caninervis* may have undergone distinct evolutionary processes that have resulted in differences in their gene content and structure. In the future, sequencing the V sex chromosome of *S. caninervis* and the U chromosome of *S. ruralis*, along with sex chromosomes of other *Syntrichia* species, will be useful for comparative purposes.

Large size and low gene density appear to be common in both U and V bryophyte sex chromosomes, for those species that have them, and high numbers of repetitive elements such as TEs are responsible for this pattern (Silva et al., 2020; Carey et al., 2021; Bowman et al., 2017; Yu et al., 2022; Gao et al., 2023). In *S. ruralis,* the sex chromosome consisted of 64.7% TEs, where autosomes had more than 50% TE content. TEs are involved in modulating plant stress regulatory networks in some plants and the large number of TEs on the sex chromosomes may be related to sex-specific differences in stress tolerance (Deneweth et al., 2022). Indeed, male and female gametophytes of *S. caninervis* have been shown to differ in several axes of stress tolerance (Stark & McLetchie, 2006; Stark et al., 2009; Stark et al., 2005a; Stark et al., 2005b).

### Uncovering the transcriptomic response to desiccation and rehydration in *S. ruralis*

To investigate the potential molecular mechanism of desiccation tolerance of *S. ruralis*, a transcriptomic analysis was conducted under a variety of conditions (**Table S8**) including under slow drying (**as defined by SD**) and rehydration following desiccation (**as defined by SDR**). An average of 16 million reads were sequenced per replicate (n = 3) per condition. Following QC, read mapping, and assigning reads to features (see Methods and **Table S8**), a single replicate was removed due to poor clustering via both principal component analysis and pairwise Euclidean distance (**Figure S4**). It is worth noting that alterations in transcript abundance, in particular during drying, may not indicate an increase in the transcription rate of the associated gene. Earlier work demonstrated that in *S. ruralis* during slow drying mRNAs were stabilized in messenger RNA particles (mRNPs) during dehydration (Wood and Oliver, 1999; Oliver 1991), which manifested as an increase in transcript abundance as visualized in northern blot analyses (Scott and Oliver 1994). Indeed, these protective mRNPs may reflect RNAs recruited into stress granules, translationally inactive cytoplasmic foci composed of RNA, protein, and metabolites that form in response to stress conditions (Kearly et al., 2022). As observed increases in transcript abundance may reflect changes in RNA stability rather than gene expression, we use the more general term “abundance” rather than “expression” below.

A series of pairwise comparisons were then performed to examine the *S. ruralis* response to dehydration and rehydration. In control vs. SD, SD vs. SDR, and control vs. SDR comparisons, 3045, 2746, and 4867 genes showing differential transcript abundance were identified, respectively (adjusted P-value < 0.001, |log2 fold change| > 2). Specifically, when comparing SD to control, transcripts for 1648 genes increased and transcripts for 1397 genes decreased in RNA abundance (**Figure 3A**). When comparing SDR to SD, transcripts for 1170 genes increased whereas 1576 genes decreased in RNA abundance (**Figure 3A**). While most of these genes were context-specific in their abundance change, a subset showed contrasting abundance patterns when comparing the transition from Control to SD and then to SDR (**Figure 3B**). Transcripts representing 340 genes increased in abundance in SD but decreased in abundance in SDR. In contrast, transcripts for 120 genes were increased in RNA abundance in both control vs. SD and SD vs. SDR comparisons, indicating that these genes may be necessary for both the dehydration and rehydration responses in *S.ruralis*. GO enrichment analysis was performed on genes showing differential RNA abundance in the control -> SD -> SDR transition. Genes with increased RNA abundance in SD (compared with control) were enriched in the GO terms: *response to abscisic acid*, *response to cold*, r*esponse to water deprivation*, *response to wounding,* and *response to karrikin* (**Figure 3C**). Interestingly, the *response to ABA* GO term was enriched in transcripts with significantly reduced abundance in rehydration, suggesting that these plants were no longer perceiving stress. The top five enriched GO terms for genes with reduced transcript abundance in SD vs. control were *photosynthesis*, *response to salicylic acid*, light stimuli (two terms), and *protein autophosphorylation* (**Figure 3C**). GO-enriched terms of genes showing decreased transcript abundance in SDR were *response to abscisic acid*, *plant-type secondary cell wall biogenesis*, *cellular response to freezing*, *response to wounding*, and *defense response to bacterium*, in which a majority of genes were associated with stress response (**Figure 3C**).

**Figure 3.**
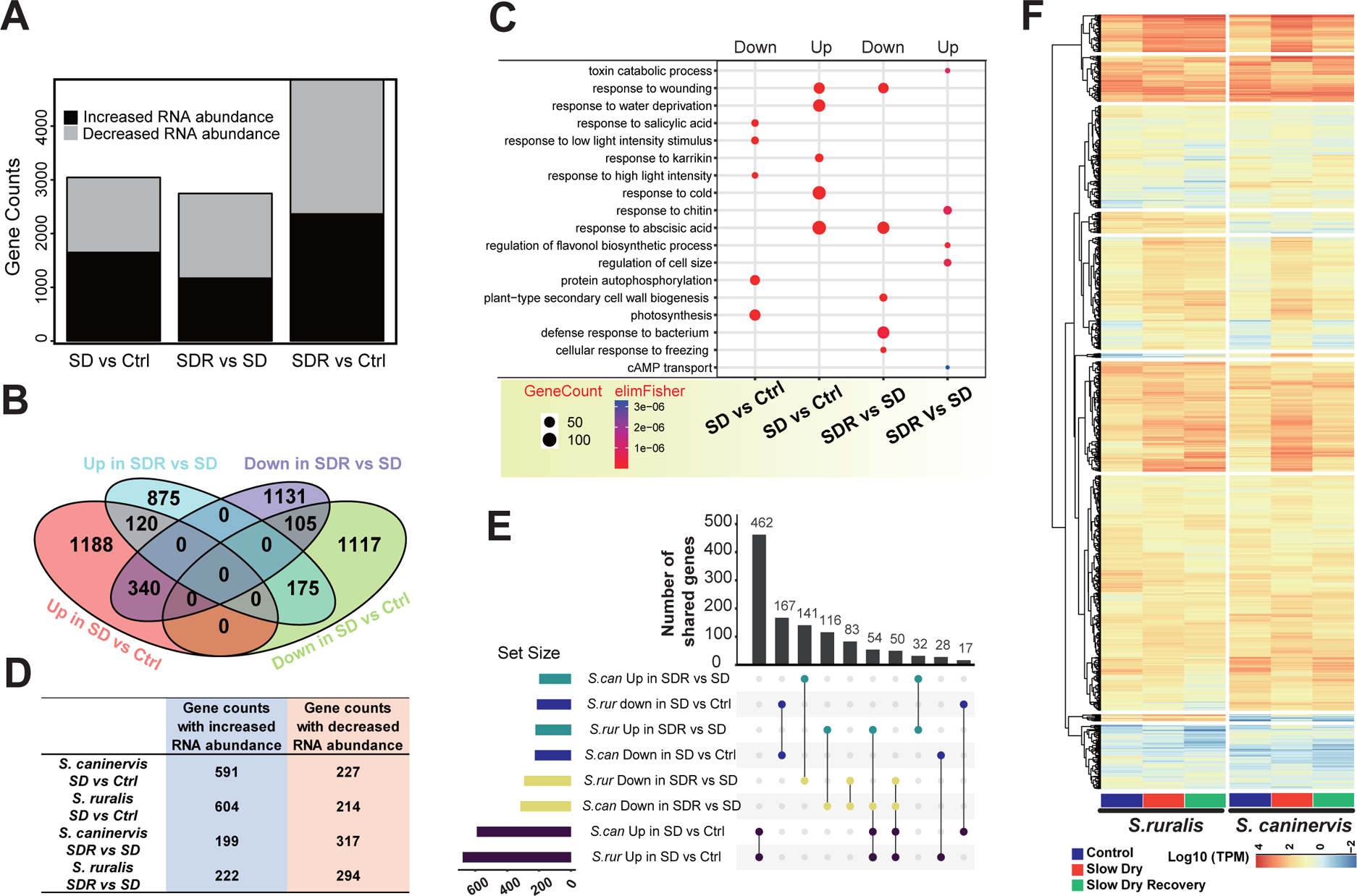
Genes that display a change in transcript abundance in response to dehydration and rehydration. **A.** The number of S. ruralis genes showing differential transcript abundance between two treatments according to a fold expression cutoff of ≥ 2 and an FDR ≤ 0.001. SD = slow dry, SDR = rehydration after the slow dry treatment. **B.** Venn diagram illustrating the number of shared and unique genes showing differential transcript abundance across different treatments. Ctrl = control. **C.** GO enrichment analysis of genes showing differential RNA abundance in Slow dry (SD) vs Ctrl (Control) or Rehydration (SDR) vs SD. The top 5 most enriched GO terms are displayed. **D.** Number of *S. ruralis* and *S. caninervis* homologous gene pairs with differential RNA abundance between SD vs Ctrl or SDR vs SD in both species. **E**. Upset plot showing homologous genes with differential RNA abundance in SD vs Ctrl or SDR vs SD of two species. **F.** Heatmap of TPMs of homologous gene pairs under SD vs Ctrl or SDR vs SD conditions.

The relatively recent divergence time between *S. ruralis* and *S. caninervis* offers a unique opportunity to search for conserved—and unique—transcriptional responses to desiccation and rehydration. To this end, publicly available RNA-sequencing data from *S. caninervis*, gathered from plants exposed to a similar set of desiccation and rehydration conditions (Silva et al., 2021), were mapped to the *S. caninervis* genome. Following similar QC filtering as performed in *S. ruralis*, changes in RNA abundance were compared for syntelogs (i.e., both collinear and sequence conserved) between these two species. Syntelogs were identified for 13,658 of the 21,169 gene models in the *S. ruralis* nuclear genome, representing 64.5%, and 82.5%, of annotated genes in *S. ruralis* and *S. caninervis*, respectively. There were 818 and 516 syntelogs showing differential RNA abundance in SD vs. control and SDR vs. SD of both *S.ruralis* and *S.caninervis*, respectively (**Figure 3D**). In the SD vs. control comparison, a majority of syntelogs had similar patterns, with RNA abundance either increasing or decreasing similarly in both species (629/818; **Figure 3E**). The analysis of the *S. caninervis* genome and transcriptome identified four dehydration-responsive transcriptomic “hot spots”, containing sets of genes that exhibited similar elevated expression patterns in response to dehydration (Silva et al., 2021). Syntelogs of these four regions were identified in the *S. ruralis* genome, however only two regions, those containing arrays of a membrane protein and a rubredoxin-like protein, increased in a similar manner as in *S. caninervis* (**Table S9**). In the SDR vs. SD comparison, a majority of syntelogs (401/516) showed different RNA abundance patterns in the two species. Interestingly, Fifty-four syntelogs exhibited increased RNA abundance in SD compared to Ctrl in both species, but *S. caninervis* showed decreased RNA abundance and *S. ruralis* displayed increased RNA abundance in SDR vs. SD comparison (**Figure 3E & Figure 3F**). These data suggest that *S. ruralis* and *S. caninervis* utilize similar molecular mechanisms to respond to dehydration, but then regulate their response to rehydration quite differently.

### Comparative genomic and transcriptomic analysis of the LEA and ELIP protein families

Certain gene families, such as late embryogenesis abundant (LEA) and early light-inducible proteins (ELIPs), have been associated with desiccation tolerance (DT) in a number of species. To better explore the evolution of these two important gene families in *S. ruralis*, candidate LEA and ELIP genes were first identified by protein sequence similarity using HMMER (Johnson et al., 2010). A total of 59 LEA genes were identified in *S. ruralis*, clustering into seven of the eight characterized LEA subgroups based on sequence and structural motifs (**Figure S5**). No LEA-6 was identified in *S. ruralis*, nor was one identified in *S. caninervis* (**Figure 4A**; Silva et al., 2021). A multiple sequence alignment of all *S. ruralis* and *S. caninervis* LEA protein sequences was used as input for inferring evolutionary relationships in this large gene family. The *S. ruralis* LEA-2 subgroup was the largest (25 genes), followed by LEA-4 (n = 17), and LEA-5 (n = 10; **Figure 4A**). Multiple instances of species-specific gene duplication and then loss were evident in these larger subgroups, as inferred by an extant duplicate in one species adjacent to a node uniting a LEA from both species (asterisks, **Figure 4A**). The ELIP gene family in *S. ruralis* is similarly large, comprising 30 members divided into four classes (I-IV; **Figure S6**). As with the LEAs, both *S. ruralis* and *S. caninervis* have experienced lineage specific duplication and loss events in the ELIP gene family. In addition, both species have experienced lineage-specific expansions that appear to have occurred through local tandem duplication events (**Figure 4B**). For instance, 9 of 30 *S. ruralis* ELIP genes are the product of tandem duplications, highlighting the volatility in these desiccation-associated gene families.

**Figure 4.**
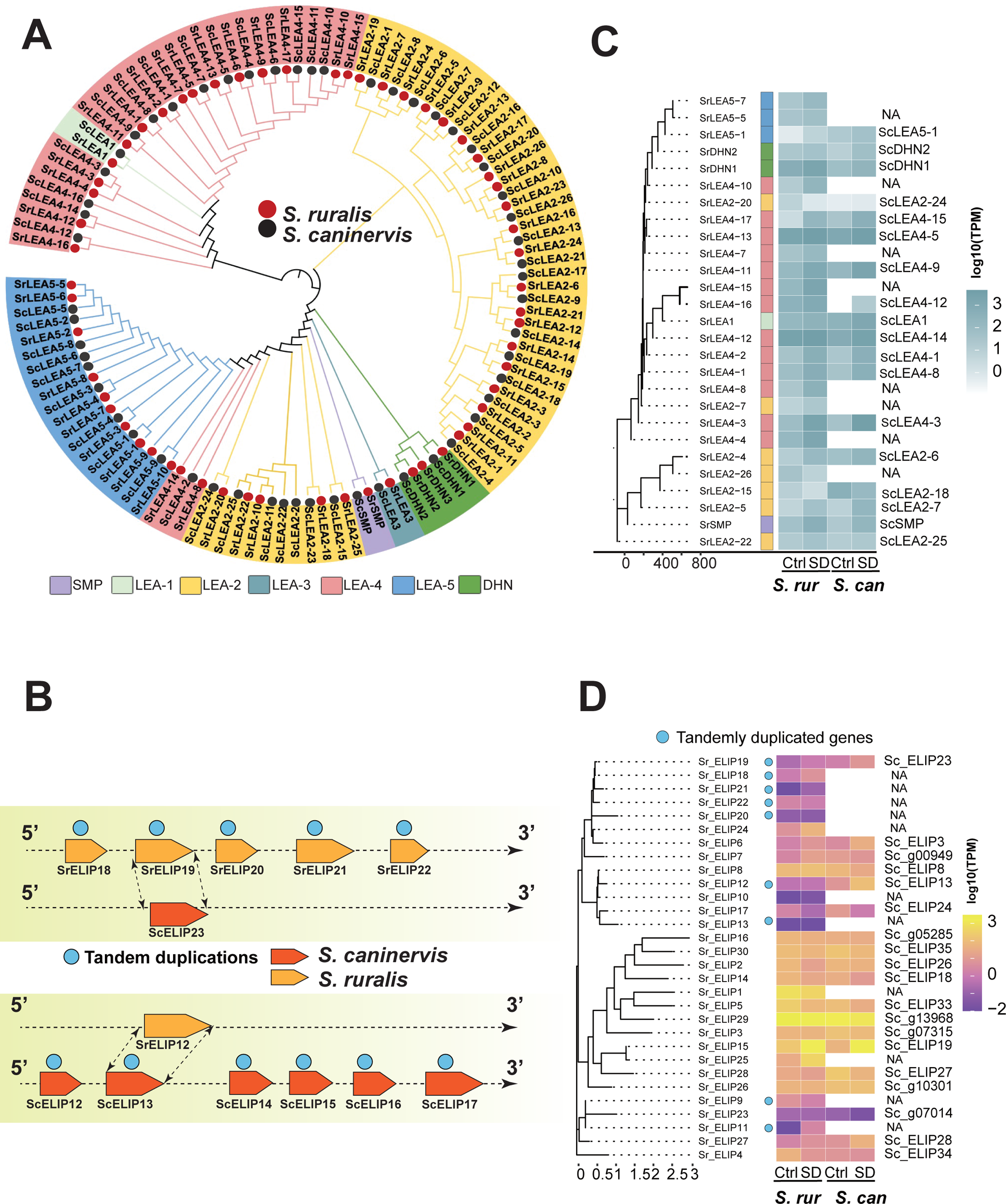
Comparison of LEA (Late embryogenesis abundant) and ELIP (Early light-induced protein) genes in *S. ruralis* and *S. caninervis*. **A.** Phylogenetic relationship between LEA gene families in *S. ruralis* and *S. caninervis*. A maximum likelihood tree was generated based on a protein multiple sequence alignment. **B.** The graph illustrated the occurrence of species-specific tandem duplication events. **C.** Heatmap of the differentially abundant *S. ruralis* LEA genes and their *S. caninervis* orthologs, where present, under control (Ctrl) and slow dry (SD) conditions. NA implies that a syntenic ortholog was not found in the *S. caninervis* genome. Normalized (transcript per million, TPM) values are shown. **D.** Heatmap of the differentially abundant *S. ruralis* ELIP genes and their *S. caninervis* orthologs, where present, under control and slow dry conditions. *Some homologous sequences where identified at syntenic locations in *S. caninervis* but were not predicted to be ELIPs based on presence of protein functional domains.

Transcriptomic analyses revealed how duplication (and loss) within these two gene families have impacted gene expression. Of the 59 identified *S. ruralis* LEA genes, transcripts of 27 of them were differentially abundant in SD conditions (**Figure 4C; Table S10**). Of these 27, nine did not have an *S. caninervis* ortholog. Of the remainder, 12 had differentially abundant transcripts for an ortholog. A similar pattern was observed for the *S. ruralis* ELIP gene family, where 11/30 ELIP transcripts were differentially abundant under desiccation conditions (six up, five down; **Figure 4D; Table S11**). Of these, three had *S. caninervis* orthologs whose transcripts were differentially abundant with similar changes in abundance in the two species. In sum, these two gene families are evolutionarily quite dynamic and look to be heavily (and differently) utilized during desiccation in these two species.

### Uncovering putative transcription factors regulating the desiccation response in *S. ruralis*

There are a number of reports discussing the similarities in which plants respond to, and transcriptionally regulate, developmental and environmental events such as embryogenesis, germination, and desiccation. As both relatives of seed plants and emerging models for desiccation tolerance, *S. ruralis* and *S. caninervis* serve as excellent systems to examine conserved regulatory mechanisms linked to water deprivation. `OrthoFinder2` was used for gene family classification and orthology prediction in 12 plant species including *S. ruralis* (**Figure S7**). Included in these 12 species are several with and without reported desiccation tolerance to help in the identification of conserved regulatory aspects of this pathway. Of the 349,650 input genes, 304,670 (87%) were assigned to 29,809 orthogroups (**Table S12**). Out of the 21,169 input nuclear genes, 19,011 (89.8%) were placed into orthogroups, with 683 of those genes falling into 195 species-specific orthogroups (not present in *S. caninervis*). *S.ruralis* has more species-specific orthogroups, compared with *S.caninervis* with 205 genes in 56 species-specific orthogroups.

To predict putative transcription factors in *S. ruralis*, the amino acid sequences of all nuclear-encoded genes were scanned using `PlantTFDB` (Tian et al., 2020), resulting in a total of 636 TFs associated with 53 families (**Table S13**). These numbers are slightly higher than those previously identified in *S. caninervis* (542 in 50 TF families) and substantially lower than in *P. patens* (1,136 in 53 families; **Table S13**; Silva et al., 2021). When comparing TF family composition between *S. ruralis* and *S. caninervis*, the B3 and C2H2 families were expanded (although not significantly) in *S. ruralis*, whereas there were no families that had experienced a significant loss.

These *S. ruralis* TFs were then assessed for a possible role in the response to desiccation by examining their change in abundance between SD and control conditions. A total of 126 (out of 636) TFs showed differential RNA abundance, with RNA levels increasing for 83 and decreasing for 53 (**Figure 5A**). Ninety-four *S. ruralis* TFs (out of 126) have syntelogs in *S. caninervis*, of which 40 also showed differential transcript abundance in SD conditions (**Figure 5A; Table S14**). Interestingly, 32 differentially abundant *S. ruralis* TFs had no identifiable syntelog in *S. caninervis*, suggesting they may impart a species-specific response to desiccation (**Figure 5A**).

**Figure 5.**
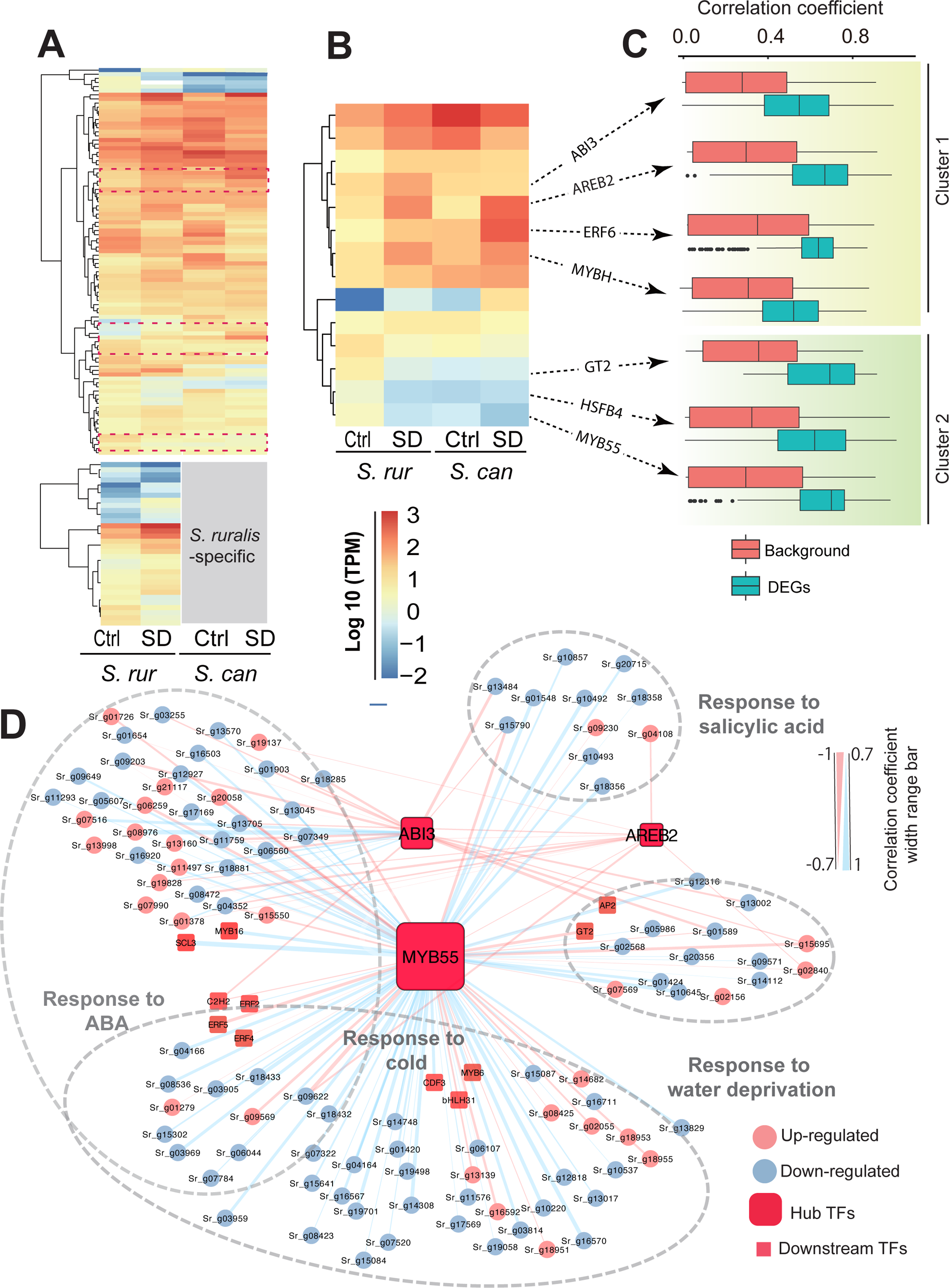
Putative TF regulators in response to dehydration (slow dry) in *S.ruralis*. **A.** The expression heatmap of putative *S.ruralis* TFs and syntenic pairs in *S. caninervis* in SD and control conditions. **B.** The expression heatmap of 14 hub TFs and *S. caninervis* syntenic pairs. **C.** Expression correlation between differentially abundant TFs and putative target genes (correlation > 0.5) and randomly selected genes (background). **C.** Regulatory network of hub TFs and putative downstream targets.

To investigate the regulation of downstream dehydration-responsive genes by these 126 TFs, the 2 Kb upstream region (promoter region) of the 3,045 genes found to be differentially abundant between control and SD conditions was extracted. Putative transcription factor binding sites were identified in these regions using AME (Analysis of Motif Enrichment; McLeay RC et al., 2010) with TF-DNA binding data from the Arabidopsis DAP motif database (O’Malley et al., 2016). From the 3,045 input genes, 121 overrepresented (enriched; p-value < 1E-60) motifs, corresponding to 120 *S. ruralis* TFs were selected for further analysis (**Table S15**). As expected based on the water limited conditions, the enriched motifs were predicted to predominantly be associated with the APETALA2/ETHYLENE RESPONSE FACTOR (AP2/ERF) and AP2/ETHYLENE RESPONSE FACTOR BINDING PROTEIN (AP2/ERFBP) families of TFs, which are well known regulators of plant abiotic stress responses. Of these 120 TFs, 14 corresponded to those that changed abundance in SD conditions (**Figures 5A** and **5B**), suggesting they may both be responding to and regulating the desiccation response in *S. ruralis*.

To assess their potential regulatory role, pairwise abundance correlation was calculated between these 14 *S. ruralis* TFs and their putative target genes across all six experimental conditions (*n* = 17 data points) and compared to a background set of genes. Seven of the fourteen TFs had a median correlation above |0.5|, with seven of these TFs significantly more correlated with their target genes than background (**Figure 5C; Figure S8**). Six of these *S. ruralis* TFs are homologous (based on amino acid sequence and domain similarities) to Arabidopsis TFs that have reported functions in drought tolerance in vegetative tissues (ABA Responsive Element Binding factor 2 [AREB2]/Sr_g16458), cell division during stress (GT2/Sr_g03037; HSFB4/Sr_g07011; MYBH/Sr_g09762; ERF6/Sr_g20827), and control gene expression during late embryogenesis - in particular the LEA gene family (ABA Insensitive 3 [ABI3]/Sr_g11739). Given their annotated functions in Arabidopsis, SrABI3 and SrAREB2 are more likely to regulate ABA responsive genes in *S. ruralis*. Several dehydration-responsive genes, including 5 LEAs, are among the 480 differentially abundant genes that contain ABI3 TF binding motifs and are correlated with SrABI3 abundance (**Figure 5D and Table S16**). Of the 346 SD-responsive genes with an AREB2 TF binding site and expression correlation > |0.5|, 29 are annotated as ABA-responsive genes based on sequence or domain similarity (**Table S16**).

### Identification of an uncharacterized regulator of dehydration in land plants

The remaining TF (Sr_g19809) is an R2R3 MYB transcription factor, a large family of regulators (both activators and repressors) well known for their diverse roles in development, metabolism, and stress responses in plants. In *S. ruralis*, *Sr_g19809* was downregulated during SD, and, due to correlation and TF binding site enrichment, appears to regulate SrGT2, SrHSFB4, and SrAREB2. Indeed, of the seven hub TFs, *Sr_g19809* appears to have the most interactions, with most of these being anti-correlated (**Figure 5D**), suggesting it is a transcriptional repressor. In addition, *Sr_g19809* is also anti-correlated with the *S. ruralis* homolog of *A. thaliana* ABI3/FUS3 (AT3G24650 and AT3G26790, respectively), which are also known to play a role in desiccation tolerance during embryogenesis (**Figure S9**; Gonzales-Morales et al., 2016).

Given the repeated incorporation of specific components of the desiccation tolerance pathway across land plants (e.g., ABI3, LEAs, and ELIPs; Marks et al., 2021), we investigated whether there were any MYB TFs similar to *Sr_g19809* in Arabidopsis. Using publicly available transcriptomic data, including samples associated with seed development, ABA application, and drought treatment (Klepikova et al., 2016; Zhao et al., 2018) we searched for *A. thaliana* MYB transcription factors anti-correlated in abundance with either AtFUS3 or AtABI3. This search uncovered a previously uncharacterized MYB TF, *AtMYB55* (AT4G01680). AtMYB55 has a similar R2R3 MYB binding domain as *Sr_g19809* (hereafter referred to as SrMYB55), is expressed during embryogenesis and germination, and shows strong negative correlation with FUS3 under drought and ABA treatment (**Figures S10A and B**), but shows a strong negative correlation with ABI3 during seed development (**Figure S10C**). An investigation of *A. thaliana* Cistrome data (O’Malley et al., 2016) identified a MYB55 binding site upstream of both ABI3 and FUS3, suggesting AtMYB55 may be a direct negative regulator of both of these genes (**Figure S10D**).

AtMYB55 is part of a clade of four similarly structured R2R3 MYB TFs in *A. thaliana* (Stracke et al., 2001). To determine the specificity of the AtMYB55-FUS3/ABI3 regulatory interaction, we also examined the closest expressed homolog of AtMYB55, AtMYB50 (AT1G57560), for its transcript abundance profile and potential ability to interact with ABI3 or FUS3. In contrast to AtMYB55, AtMYB50 was not correlated with ABI3 and FUS3 (**Figure S10A-C**), nor is a MYB50 binding site seen in the promoter elements of these two genes. In addition, AtMYB50 is induced by the biotic-stress associated phytohormones JA and SA (Katiyar et al., 2012) and thus likely works to regulate a distinct set of genes from AtMYB55.

Given that SrMYB55 regulates a number of ABA-associated genes and AtMYB55 is expressed during germination (**Figure S10D**), we hypothesized that AtMYB55 mutants might display a sensitivity to ABA at this developmental stage. We identified a T-DNA insertion line within the third exon of AtMYB55 (**Figure 6A**). Both wild type and *Atmyb55* showed 100 % germination after 24 hours on plates without ABA (**Figure 6B**). On the plates with 0.1 uM, 0.3 uM, and 0.5 uM ABA, wild type seeds displayed delayed germination, but germinated at levels close to control plates after 72 hours. In contrast, the *Atmyb55* background showed dramatically delayed germination, and at the maximum ABA concentration (0.5 uM), never reached control levels of germination (**Figure 6C**). No variation in number or density of lateral roots, nor in length of primary root, was observed between wild type and *Atmyb55*, suggesting the ABA sensitivity in this background is restricted to germination under normal growth conditions (**Figure S11**). In sum, MYB55 appears to be negatively regulating ABI3/FUS3 in both *A. thaliana* and *S. ruralis*.

**Figure 6.**
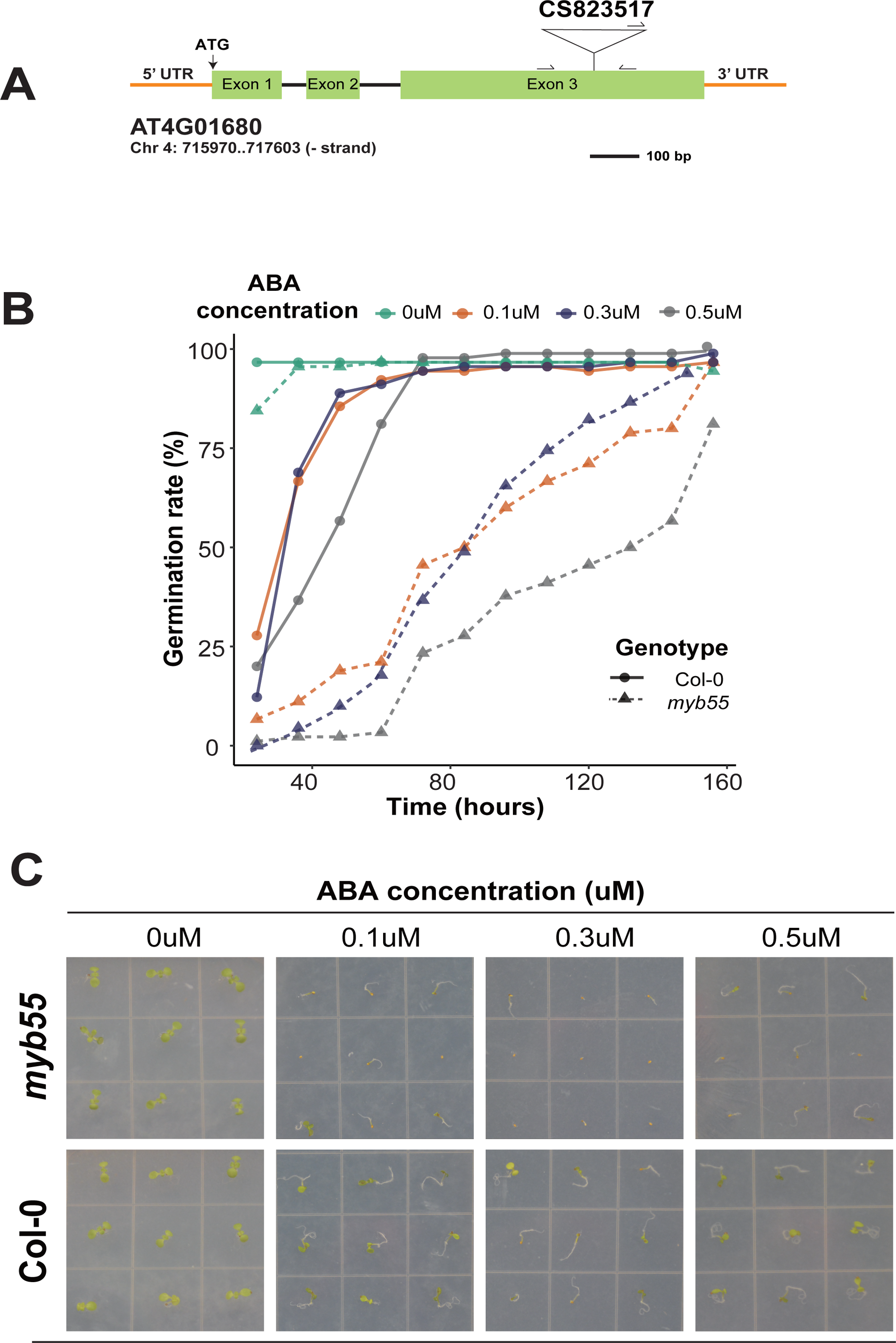
*MYB55* is sensitive to ABA in Arabidopsis. **A.** Schematic diagram of the T-DNA insertion of the *MYB55* gene. **B.** Seed germination rate of *myb55* and Col-0 lines in response to different concentrations of ABA. Numbers of germinated seedlings were recorded from 0 to 156 h after stratification on 1/2 MS agar plates containing 0, 0.1, 0.3 or 0.5 μM ABA. **C.** Phenotypes of seedlings grown for 7 days after stratification on agar plates containing 0, 0.1, 0.3 or 0.5 μM ABA.

MYB55 transcript abundance appears to be negatively correlated with water content in both species (**Figure 5B** and **Figures 10A-C**). A compelling model arises in which MYB55 is tightly linked to perceived water stress in an ABA-dependent manner. As water stress increases, MYB55 levels are decreased, allowing for increased levels of either ABI3 or FUS3, depending on the developmental stage. While it remains to be seen if MYB55 directly binds the combined ABI3-FUS3 homolog in *S. ruralis*, the observed regulatory interactions in both species are suggestive of a conserved mechanism between *A. thaliana* and *S. ruralis*, and maybe across land plants in general. In conclusion, our work here highlights the exciting contributions that studying desiccation tolerance in *S. ruralis* can have on our understanding of abiotic stress tolerance across the land-plant lineage.

## Author Contributions and Acknowledgements

XZ, JTBE, BDM, ADLN, and MJO developed the project. ATS isolated nucleic acids for sequencing, assembled and annotated the genome. XZ, JTBE, and LY performed bioinformatic analyses. XZ and AKJ performed the Arabidopsis analyses. XZ, JTBE, ADLN, and MJO wrote the manuscript. All authors edited and approved the manuscript. We would like to acknowledge support from NSF Dimensions of Biodiversity Program grants 1638972 (to MJO) and 1638956 (to BDM), NSF IOS PGRP 2023310 (to ADLN) and NSF IOS PGRP 2102120 (to ADLN). JTBE was funded by the NSF CAREER Award 2144011. We would like to thank the Boyce Thompson Institute PGRP REU program for hosting AKJ (REU support for AKJ from NSF #2023310). We would like to thank Dr. Jingjing Zhai (Cornell University) for bioinformatic analysis support. The authors would like to thank other members of the Nelson lab for their insightful feedback during the preparation of this manuscript. Authors would like to thank members of the Oliver lab, Kate Guill and James Elde, for their technical expertise in maintaining moss cultures and the isolation of high molecular weight genomic DNA.

## Data Availability

*S. ruralis* isolates are available by request from ADLN. The *S. ruralis* genome assembly and annotation are publicly available at CoGe (genomevolution.org). DNA and RNA sequencing data are available at NCBI (Bioprojects XXX and XXX).

---

**Figure S1.** Comparison of genome size and annotated gene content for published moss genomes. References: *Syntrichia caninervis* (Silva et al., 2020), *Takakia lepidozioides* (Hu et al., 2023), *Physcomitrium patens* (Lang et al., 2018; Rensing et al., 2020), *Ceratodon purpureus*, (Carey et al., 2021), *Entodon seductrix* (Yu et al., 2022), *Fontinalis antipyretica* (Yu et al., 2020), *Calohypnum plumiforme* (Mao et al., 2020), *Pohlia nutans* (Liu et al., 2022), *Hypnum curvifolium* (Yu et al., 2022). *Bryum argenteum* (Gao et al., 2023), *Pleurozium schreberi* (Pederson et al., 2019), *Sphagnum angustifolium* and *Sphagnum divinum* (Healey et al., 2023).

**Figure S2.** Classification of transposable elements (TEs) and proportion of different classes of TEs displayed by the nested pie chart. LTR: Long Terminal Repeat; TIR: Terminal Inverted Repeat.

**Figure S3.** Large-scale genome conservation between *S. ruralis*, *S. caninervis* and *P. patens* generated by McScan. Each node (connection point) represents a genomic region, and the edges (lines) connecting the nodes represent syntenic blocks or collinear regions of syntenic genes. The length and thickness of the edges indicate the size and similarity of the syntenic blocks, respectively. Conserved syntenic blocks are depicted in shared colors. The green edges reflect a chromosomal rearrangement event (fusion event) between *S. caninervis* Chr. 2 and 8 and *S. ruralis* Chr. 11.

**Figure S4.** Quality control of *S.ruralis* samples. **A.** Pairwise Euclidean distance among *S.ruralis* samples. **B.** Principal Component Analysis (PCA) of *S.ruralis* samples. **C.** Correlation between *S.ruralis* slow dry rehydrate (SDR) replicates.

**Figure S5.** Consensus patterns of LEA subgroups. **A.** Two HMM logos of DNH subgroups. **B.** The sequence alignment of *S. ruralis* DHN genes. **C.** The HMM logo of LEA 5 subgroup. **D.** The sequence alignment of *S. ruralis* LEA 5 genes. The red boxes indicated the consensus sequences.

**Figure S6.** Phylogenetic relationship between ELIP gene families in *S. ruralis* and *S. caninervis*. A maximum likelihood tree was generated based on a protein multiple sequence alignment.

**Figure S7.** A phylogeny of select land plants suggesting that desiccation tolerance has evolved multiple times in the land plant lineage. *S. ruralis*, *S. caninervis* and *O. thomaeum* are desiccation-tolerant (DT), *P. patens* and *M. polymorpha* exhibited vegetative DT (vDT) under certain environmental conditions, rest of species are desiccation-sensitive.

**Figure S8.** The density plot of correlation between putative TFs and downstream targets. TFs highlighted in bold showed the correlation with targets > 0.5 and < −0.5.

**Figure S9.** Heatmap of transcript abundance values for *S. ruralis* FUS3, ABI3, and MYB55.

**Figure S10.** Heat map of transcript abundance data for Arabidopsis MYB55, MYB50, LEC1, ABI3, and FUS3 during drought and ABA treatment, as well as during seed development and germination.

**Figure S11.** Phototypes of *myb55* and Col-0 seedings that germinated on plates with 0 μM ABA and transplanted and grown for a week on agar plates containing 0.3, 0.5, 1 and 2 μM ABA.

## References

Alexa, A. and Rahnenführer, J., 2009. Gene set enrichment analysis with topGO. Bioconductor Improv, 27, pp.1–26.

Belnap, J. and O. L. Lange. 2003. Structure and functioning of biological soil crusts: a synthesis. pp. 471–479 in J. Belnap & O. L. Lange, eds, Biological Soil Crusts: Structure, Function, and Management. Ecological Studies vol. 150. Springer-Verlag, Berlin.

Brinda, J. C., Jáuregui-Lazo, J. A., Oliver, M. J., & Mishler, B. D. (2021). Notes on the genus Syntrichia with a revised infrageneric classification and the recognition of a new genus Syntrichiadelphus (Bryophyta, Pottiaceae). Phytologia, 103(4).

Carey, S. B., Jenkins, J., Lovell, J. T., Maumus, F., Sreedasyam, A., Payton, A. C., … & McDaniel, S. F. (2021). Gene-rich UV sex chromosomes harbor conserved regulators of sexual development. Science Advances, 7(27), eabh2488.

Coe, K.K., Greenwood, J.L., Slate, M.L., Clark, T.A., Brinda, J.C., Fisher, K.M., Mishler, B.D., Bowker, M.A., Oliver, M.J., Ebrahimi, S. and Stark, L.R., 2021. Strategies of desiccation tolerance vary across life phases in the moss *Syntrichia caninervis*. American journal of botany, 108(2), pp.249–262.

Fajkus, P., Peška, V., Závodník, M., Fojtová, M., Fulnečková, J., Dobias, Š., Kilar, A., Dvořáčková, M., Zachová, D., Nečasová, I., Sims, J., Sýkorová, E., & Fajkus, J. (2019). Telomerase RNAs in land plants. Nucleic acids research, 47(18), 9842–9856. 10.1093/nar/gkz695

Fajkus P, Kilar A, Nelson ADL, Holá M, Peška V, Goffová I, Fojtová M, Zachová D, Fulnečková J, Fajkus J. Evolution of plant telomerase RNAs: farther to the past, deeper to the roots. Nucleic Acids Res. 2021 Jul 21;49(13):7680–7694. doi: 10.1093/nar/gkab545.

Gao, B., Li, X., Liang, Y., Zhang, J., Oliver, M., & Zhang, D. (2023). Characterization of UV sex chromosomes and synteny-guided phylogenomic resolution of subgenomes in Bryopsida mosses.

Gao, C.H., Yu, G. and Cai, P., 2021. ggVennDiagram: an intuitive, easy-to-use, and highly customizable R package to generate Venn diagram. Frontiers in Genetics, p.1598.

González-Morales SI, Chávez-Montes RA, Hayano-Kanashiro C, Alejo-Jacuinde G, Rico-Cambron TY, de Folter S, Herrera-Estrella L. Regulatory network analysis reveals novel regulators of seed desiccation tolerance in Arabidopsis thaliana. Proc Natl Acad Sci U S A. 2016 Aug 30;113(35):E5232–41.

Greenwood, J.L. and Stark, L.R. 2014. The rate of drying determines the extent of desiccation tolerance in *Physcomitrella patens*. Functional Plant Biology 41: 460–467.

Healey, A. L., Piatkowski, B., Lovell, J. T., Sreedasyam, A., Carey, S. B., Mamidi, S., … & Shaw, A. J. (2023). Newly identified sex chromosomes in the Sphagnum (peat moss) genome alter carbon sequestration and ecosystem dynamics. Nature Plants, 9(2), 238–254.

Jauregui-Lazo, J., Brinda, J. C., GoFlag Consortium, & Mishler, B. D. (2023). The phylogeny of Syntrichia: An ecologically diverse clade of mosses with an origin in South America. American Journal of Botany, 110(1), e16103.

Johnson, L. Steven, Sean R. Eddy, and Elon Portugaly. “Hidden Markov model speed heuristic and iterative HMM search procedure.” BMC bioinformatics 11 (2010): 1–8.

Katiyar, A., Smita, S., Lenka, S.K. et al. Genome-wide classification and expression analysis of *MYB* transcription factor families in rice and Arabidopsis. BMC Genomics 13, 544 (2012).

Klepikova, Anna V., et al. “A high resolution map of the Arabidopsis thaliana developmental transcriptome based on RNA-seq profiling.” The Plant Journal 88.6 (2016): 1058–1070.

Koster K.L., Balsamo R.A., Espinoza C. and Oliver M.J. 2010. Desiccation sensitivity and tolerance in the moss *Physcomitrella patens*: assessing limits and damage. Plant Growth Regulation 62: 293–302.

Lang, Daniel, Kristian K. Ullrich, Florent Murat, Jörg Fuchs, Jerry Jenkins, Fabian B. Haas, Mathieu Piednoel, et al. “The *Physcomitrella patens* chromosome-scale assembly reveals moss genome structure and evolution.” The Plant Journal 93, no. 3 (2018): 515–533

Lang, Daniel, Kristian K. Ullrich, Florent Murat, Jörg Fuchs, Jerry Jenkins, Fabian B. Haas, Mathieu Piednoel, et al. “The Physcomitrella patens chromosome-scale assembly reveals moss genome structure and evolution.” The Plant Journal 93, no. 3 (2018): 515–533.

Li, X., Yang, R., Liang, Y., Gao, B., Li, S., Bai, W., Oliver, M.J. and Zhang, D., 2023. The ScAPD1-like gene from the desert moss *Syntrichia caninervis* enhances resistance to *Verticillium dahliae* via phenylpropanoid gene regulation. The Plant Journal, 113(1), 75–91

Liu, S., Fang, S., Cong, B., Li, T., Yi, D., Zhang, Z., … & Zhang, P. (2022). The Antarctic moss Pohlia nutans genome provides insights into the evolution of bryophytes and the adaptation to extreme terrestrial habitats. Frontiers in plant science, 13, 920138.

Love, Michael I., Wolfgang Huber, and Simon Anders. “Moderated estimation of fold change and dispersion for RNA-seq data with DESeq2.” Genome biology 15, no. 12 (2014): 1–21.

Lyons, Eric, and Michael Freeling. “How to usefully compare homologous plant genes and chromosomes as DNA sequences.” The Plant Journal 53, no. 4 (2008): 661–673.

Marks RA, Farrant JM, Nicholas McLetchie D, VanBuren R. Unexplored dimensions of variability in vegetative desiccation tolerance. Am J Bot. 2021 Feb;108(2):346–358.

McLeay, Robert C., and Timothy L. Bailey. “Motif Enrichment Analysis: a unified framework and an evaluation on ChIP data.” BMC bioinformatics 11, no. 1 (2010): 1–11

Mishler, BD and Oliver, MJ. (2009). Putting Physcomitrella on the tree of life: the evolution and ecology of mosses. In: The Moss Physcomitrella patens. Ed. C. Knight, P-F Perroud and D Cove. Annual Plant Reviews. Wiley Blackwell.

Naramoto, S., Hata, Y., Fujita, T. and Kyozuka, J., 2022. The bryophytes *Physcomitrium patens* and *Marchantia polymorpha* as model systems for studying evolutionary cell and developmental biology in plants. The Plant Cell, 34(1), pp.228–246.

O’Leary, Nuala A., Mathew W. Wright, J. Rodney Brister, Stacy Ciufo, Diana Haddad, Rich McVeigh, Bhanu Rajput et al. “Reference sequence (RefSeq) database at NCBI: current status, taxonomic expansion, and functional annotation.” Nucleic acids research 44, no. D1 (2016): D733–D745.

Oldenhof H., Wolkers W.F., Bowman J.L., Tablin F., Crowe J.H. 2006. Freezing and desiccation tolerance in the moss *Physcomitrella patens*: an in situ Fourier transform infrared spectroscopic study. Biochimica et Biophysica Acta 1760, 1226–1234

Oliver, M. J., J. Velten, and B. D. Mishler. 2005. Desiccation tolerance in bryophytes: a reflection of the primitive strategy for plant survival in dehydrating habitats? Integrative and Comparative Biology 45:788–799.

Oliver, MJ, and Bewley, JD. (1984). Plant desiccation and protein synthesis: VI. Changes in protein synthesis elicited by desiccation of the moss *Tortula ruralis* are effected at the translational level. Plant Physiology. 74: 923–927.

Oliver, MJ. (1991) Influence of protoplasmic water loss on the control of protein synthesis in the desiccation-tolerant moss *Tortula ruralis*: Ramifications for a repair-based mechanism of desiccation-tolerance. Plant Physiology. 97:1501–1511.

Oliver, MJ, Mishler, BD, and Quisenberry JE. (1993). Comparative measures of desiccation-tolerance in the *Tortula ruralis* complex. I. Variation in damage control and repair. Am. J. Bot. 80:127–136

Oliver, MJ. (2009) Biochemical and molecular mechanisms of desiccation tolerance in bryophytes. In Bryophyte Biology, 2nd Edition. Eds, Shaw, J and Goffinet, B. Cambridge Press. pp 269

Ou, Shujun, Weija Su, Yi Liao, Kapeel Chougule, Jireh RA Agda, Adam J. Hellinga, Carlos Santiago Blanco Lugo et al. “Benchmarking transposable element annotation methods for creation of a streamlined, comprehensive pipeline.” Genome biology 20, no. 1 (2019): 1–18.

O’Malley, Ronan C., Shao-shan Carol Huang, Liang Song, Mathew G. Lewsey, Anna Bartlett, Joseph R. Nery, Mary Galli, Andrea Gallavotti, and Joseph R. Ecker. “Cistrome and epicistrome features shape the regulatory DNA landscape.” Cell 165, no. 5 (2016): 1280–1292.

Proctor, M.C., Oliver, M.J., Wood, A.J., Alpert, P., Stark, L.R., Cleavitt, N.L. and Mishler, B.D., 2007. Desiccation-tolerance in bryophytes: a review. The bryologist, 110(4), pp.595–621.

Rensing, S.A., Goffinet, B., Meyberg, R., Wu, S.Z. and Bezanilla, M., 2020. The moss *Physcomitrium (Physcomitrella) patens*: a model organism for non-seed plants. The Plant Cell, 32(5), pp.1361–1376.

Reski, R., Bae, H. and Simonsen, H.T., 2018. *Physcomitrella patens*, a versatile synthetic biology chassis. Plant cell reports, 37, pp.1409–1417.

Saitox, Rintaro, et al. “A travel guide to Cytoscape plugins.” Nature methods 9.11 (2012): 1069–1076.

Schonbeck, M.W. and Bewley, J.D., 1981. Responses of the moss *Tortula ruralis* to desiccation treatments. II. Variations in desiccation tolerance. Canadian Journal of Botany, 59(12), pp.2707–2712.

Scott, HB and Oliver, MJ. (1994) Accumulation and polysomal recruitment of transcripts in response to desiccation and rehydration of the moss *Tortula ruralis*. Journal Experimental Botany 45:577–583

Silva, A.T., Gao, B., Fisher, K.M., Mishler, B.D., Ekwealor, J.T., Stark, L.R., Li, X., Zhang, D., Bowker, M.A., Brinda, J.C., Coe, K.K., and Oliver, M.J. 2021. To dry perchance to live: Insights from the genome of the desiccation-tolerant biocrust moss *Syntrichia caninervis*. The Plant Journal. 105, 1339–1356

Stark, L. R., & McLetchie, D. N. (2006). Gender-specific heat-shock tolerance of hydrated leaves in the desert moss Syntrichia caninervis. Physiologia Plantarum, 126(2), 187–195.

Stark, L. R., D. N. McLetchie, and B. D. Mishler. 2005. Sex expression, plant size, and spatial segregation of the sexes across a stress gradient in the desert moss Syntrichia caninervis. Bryologist 108: 183–193

Stark, L. R., J. C. Brinda, D. N. McLetchie & M. J. Oliver. 2012. Extended periods of hydration do not elicit dehardening to desiccation tolerance in regeneration trials of the moss *Syntrichia caninervis*. International Journal of Plant Sciences 173: 333–343

Stark, L. R., McLetchie, D. N., & Mishler, B. D. (2005). Sex expression, plant size, and spatial segregation of the sexes across a stress gradient in the desert moss Syntrichia caninervis. The Bryologist, 108(2), 183–193.

Stark, L. R., McLetchie, D. N., & Roberts, S. P. (2009). Gender differences and a new adult eukaryotic record for upper thermal tolerance in the desert moss Syntrichia caninervis. Journal of Thermal Biology, 34(3), 131–137.

Stark, L. R., Nichols II, L., McLetchie, D. N., & Bonine, M. L. (2005). Do the sexes of the desert moss Syntrichia caninervis differ in desiccation tolerance? A leaf regeneration assay. International Journal of Plant Sciences, 166(1), 21–29.

Stark, L.R. “Ecology of desiccation tolerance in bryophytes: A conceptual framework and methodology,” The Bryologist 120(2), 130–165, (2 June 2017). 10.1639/0007-2745-120.2.130

Stracke R, Werber M, Weisshaar B. The R2R3-MYB gene family in Arabidopsis thaliana. Curr Opin Plant Biol. 2001 Oct;4(5):447–56.

Tian, Feng, De-Chang Yang, Yu-Qi Meng, Jinpu Jin, and Ge Gao. “PlantRegMap: charting functional regulatory maps in plants.” Nucleic acids research 48, no. D1 (2020): D1104–D1113.

Wood, AJ, and Oliver, MJ. (1999) Translational control in plant stress: Formation of messenger ribonucleoprotein complexes (mRNPs) in Tortula ruralis in response to desiccation. The Plant Journal. 18(4):359–370

Wu, S.Z., Ryken, S.E. and Bezanilla, M., 2023. CRISPR-Cas9 Genome Editing in the Moss *Physcomitrium* (Formerly *Physcomitrella*) *patens*. Current Protocols, 3(4), p.e725.

Xiao, L., Yobi, A., Koster, K.L., He, Y., and Oliver, M.J. (2018) Desiccation tolerance in *Physcomitrella patens*: rate of dehydration and the involvement of endogenous ABA. Plant, Cell & Environment. 41: 275–284

Yu, J., Cai, Y., Zhu, Y., Zeng, Y., Dong, S., Zhang, K., … & Liu, Y. (2022). Chromosome-level genome assemblies of two Hypnales (mosses) reveal high intergeneric synteny. Genome Biology and Evolution, 14(2), evac020.

Yu, J., Li, L., Wang, S., Dong, S., Chen, Z., Patel, N., … & Liu, Y. (2020). Draft genome of the aquatic moss Fontinalis antipyretica (Fontinalaceae, Bryophyta). Gigabyte, 2020.

Zhang, J., Zhang, Y.M., Downing, A., Wu, N. and Zhang, B.C., 2011. Photosynthetic and cytological recovery on remoistening *Syntrichia caninervis* Mitt., a desiccation-tolerant moss from Northwestern China. Photosynthetica, 49, pp.13–20.

Zhao, Xinyue, et al. “Global identification of Arabidopsis lncRNAs reveals the regulation of MAF4 by a natural antisense RNA.” Nature communications 9.1 (2018): 5056.

Zhong, B., Fong, R., Collins, L. J., McLenachan, P. A., & Penny, D. (2014). Two new fern chloroplasts and decelerated evolution linked to the long generation time in tree ferns. Genome Biology and Evolution, 6(5), 1166–1173.

