## Supplementary figures and images for "*Syntrichia ruralis*: Emerging model moss genome reveals a conserved and previously unknown regulator of desiccation in flowering plants"

### Supplemental figure 1

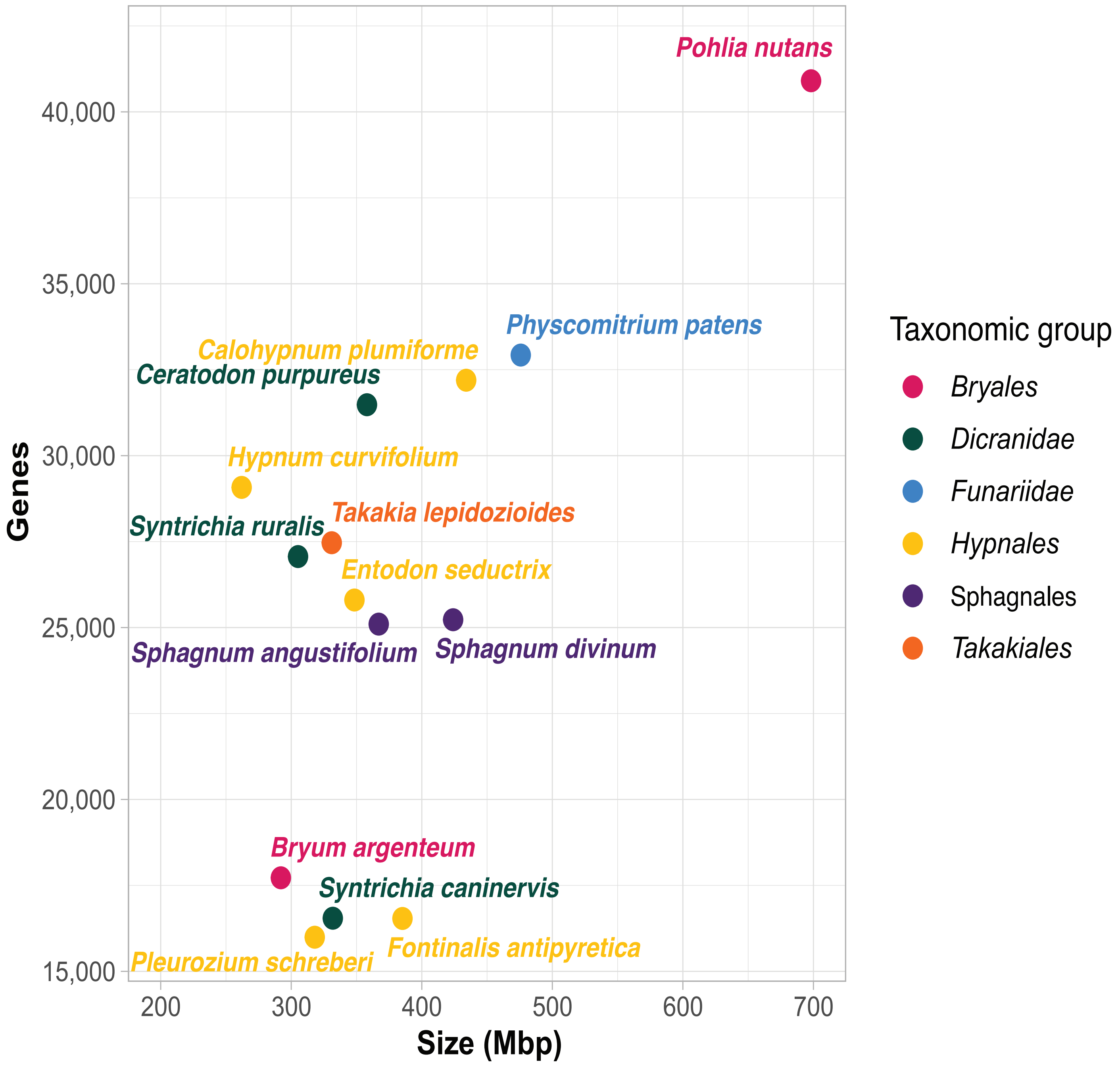

### Supplemental figure 2

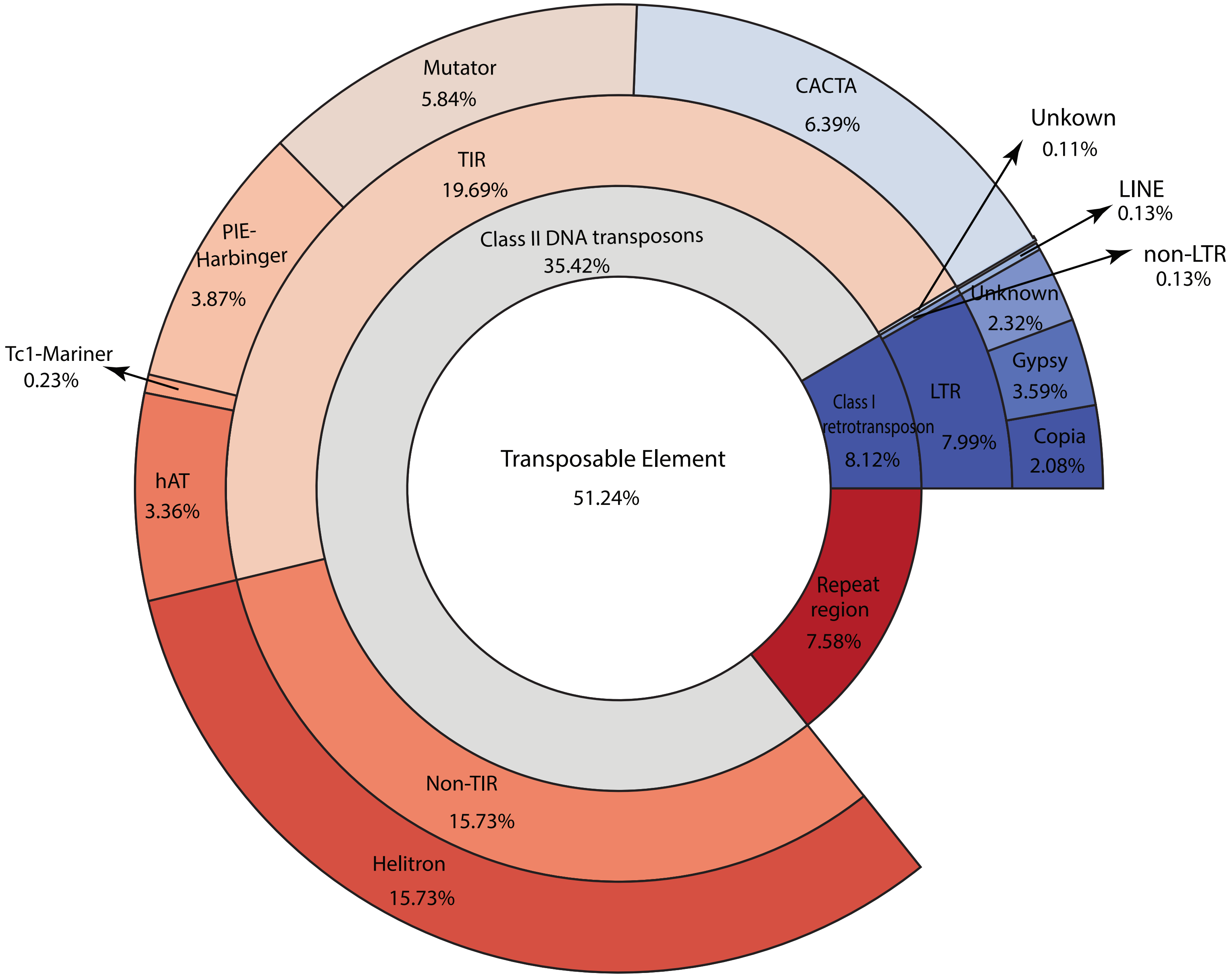

### Supplemental figure 3

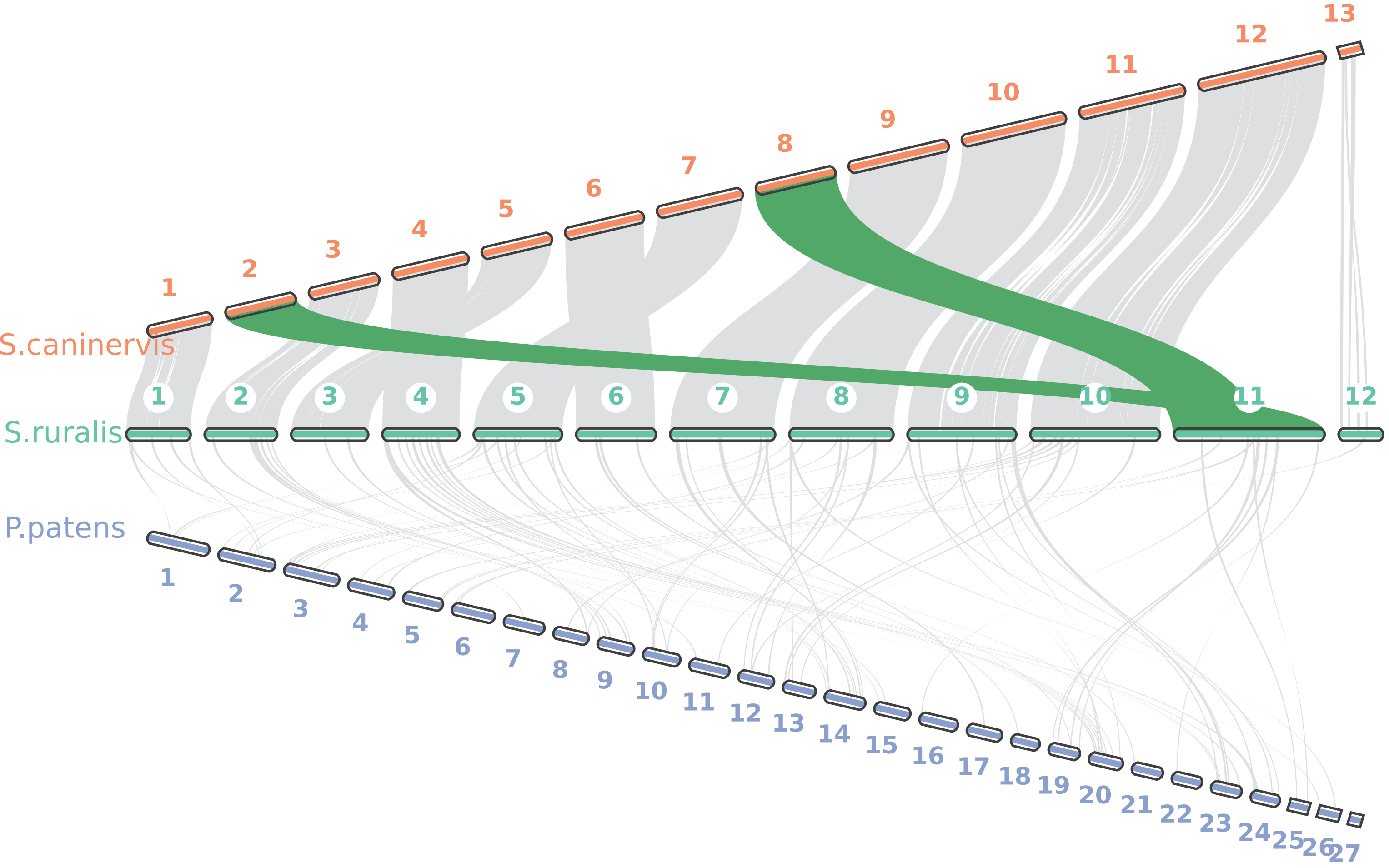

### Supplemental figure 4

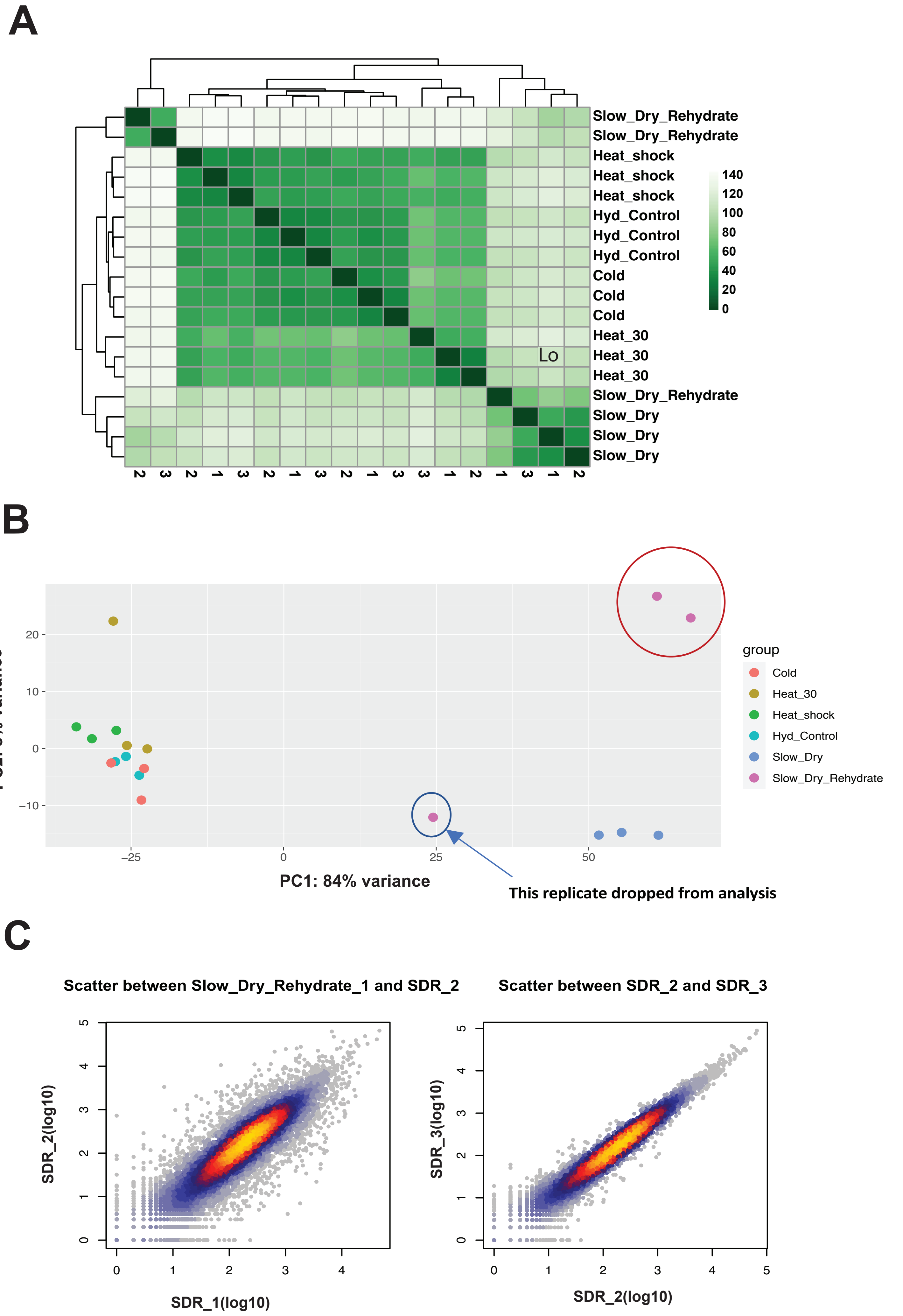

### Supplemental figure 5

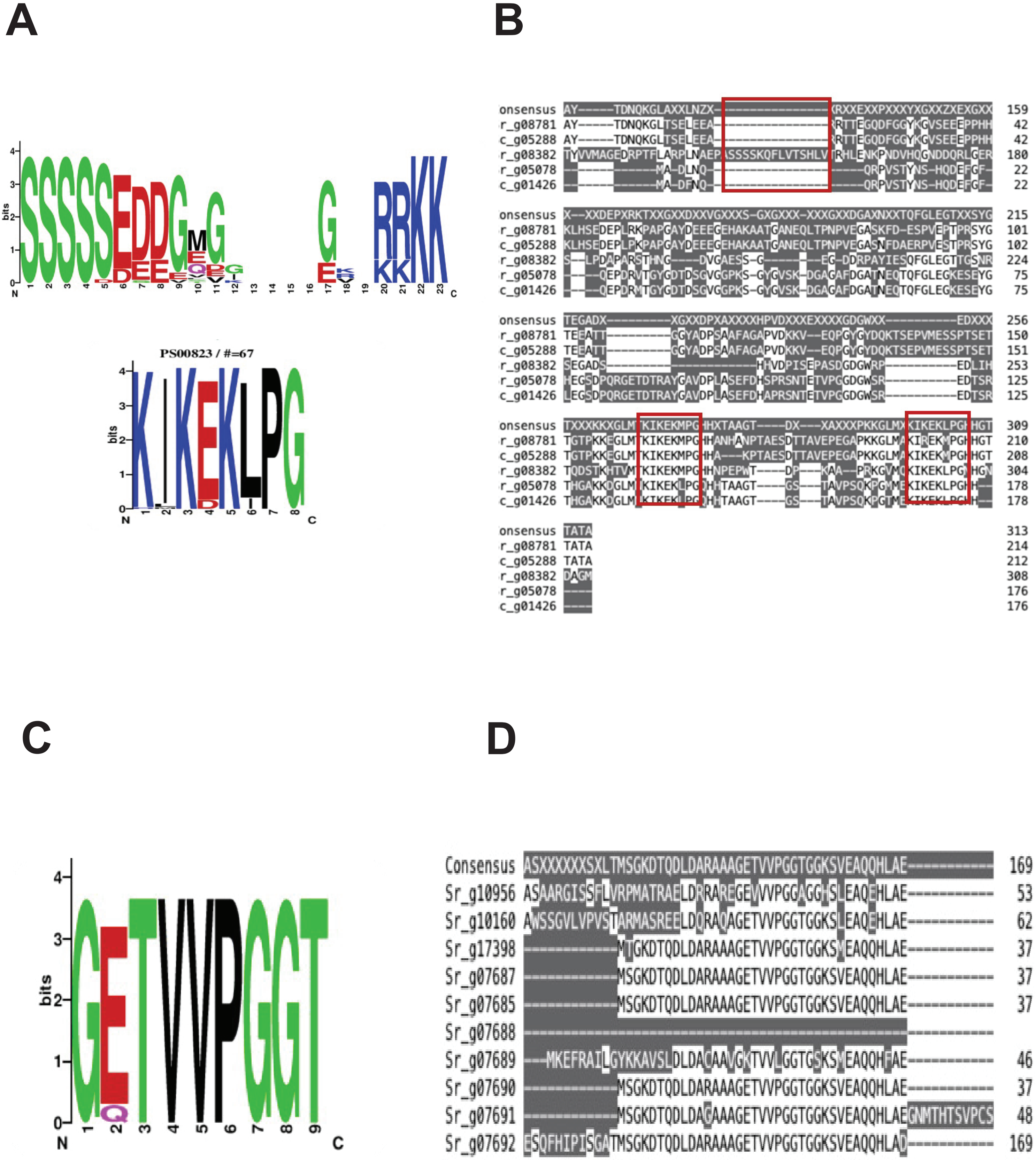

### Supplemental figure 6

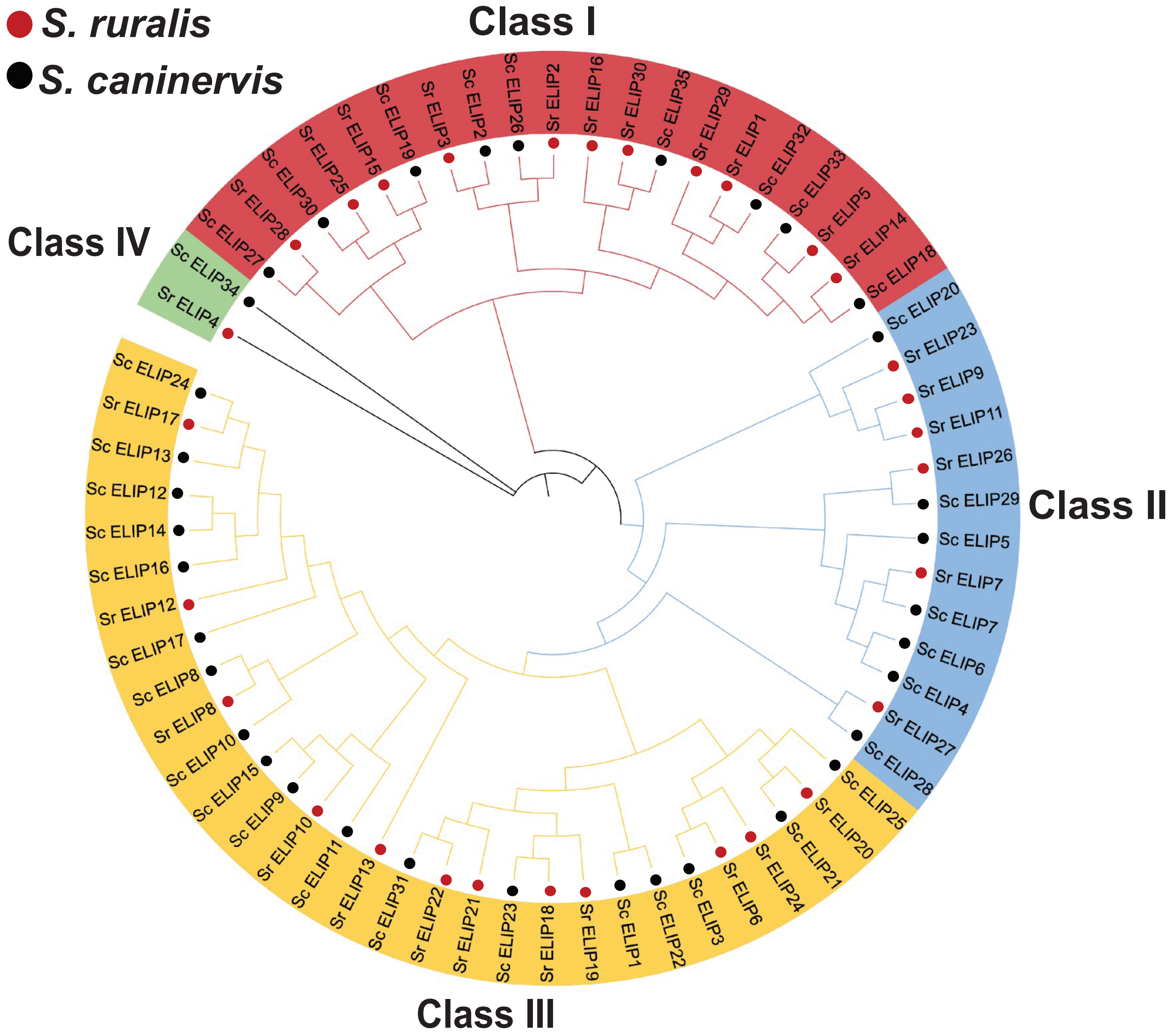

### Supplemental figure 7

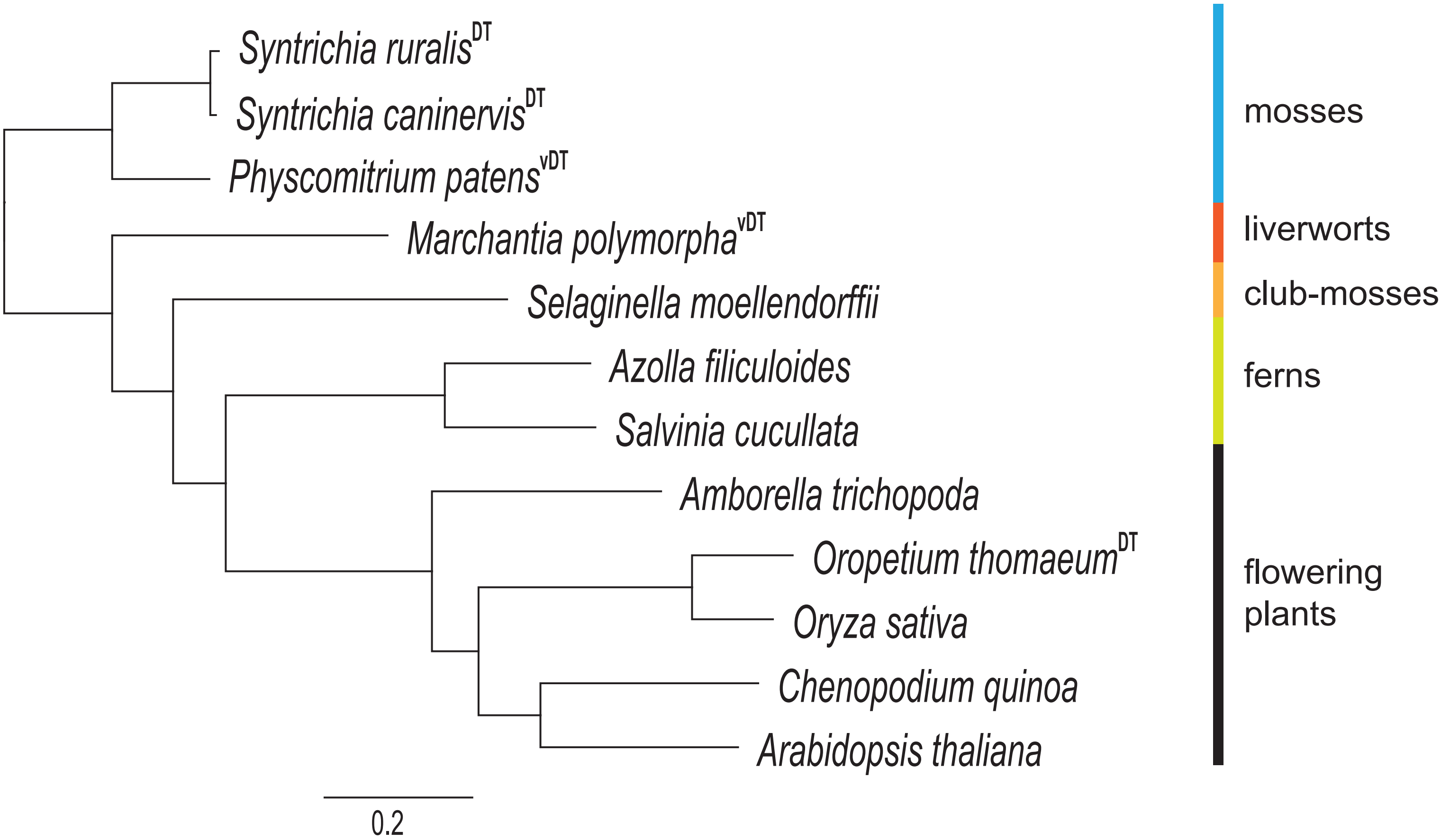

### Supplemental figure 8

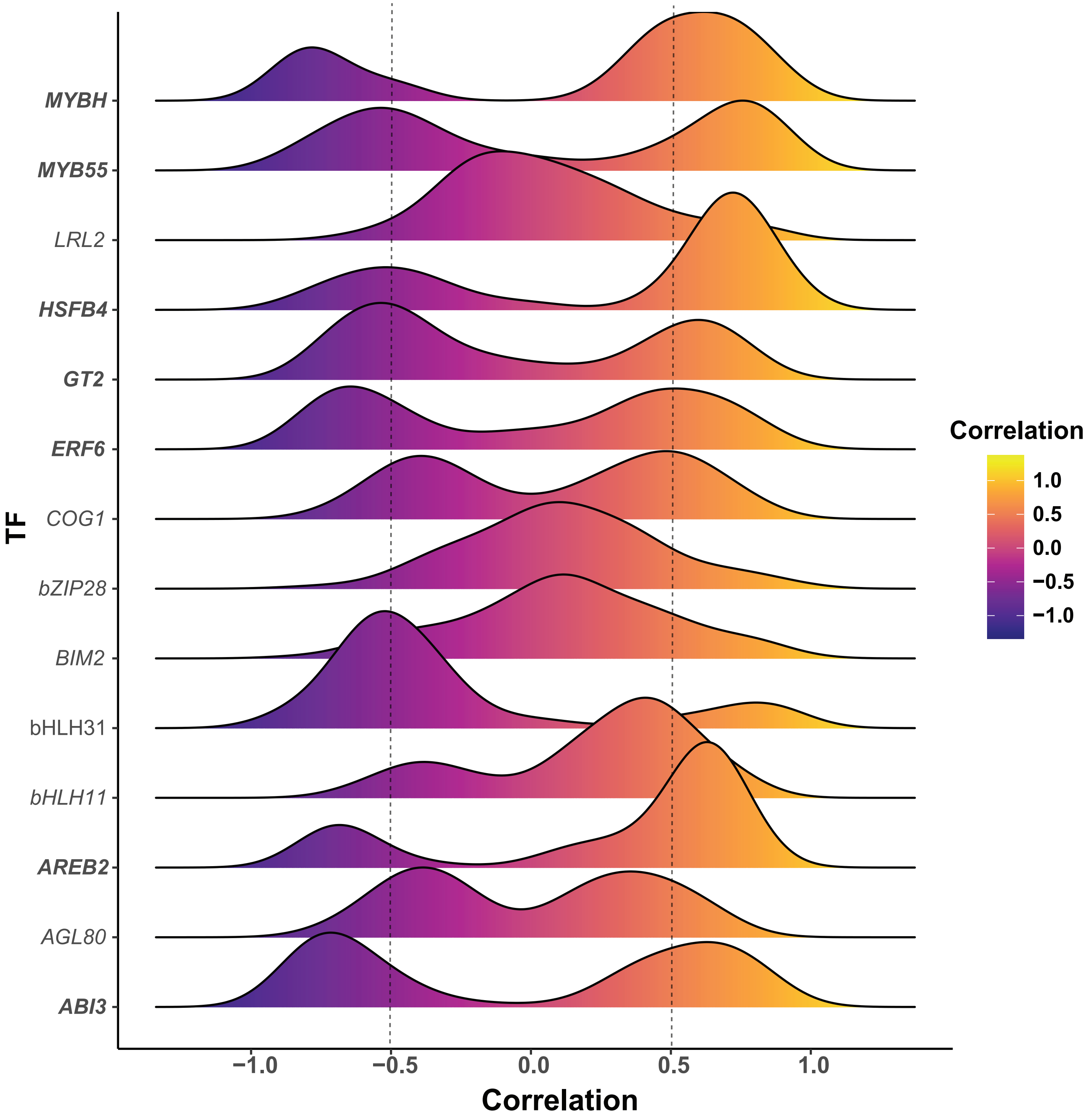

### Supplemental figure 9

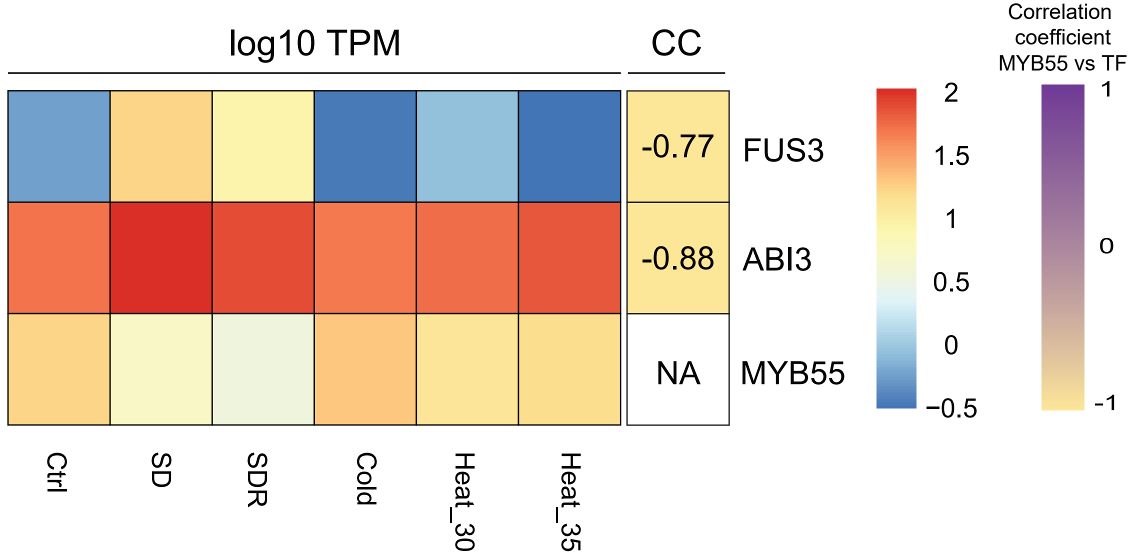

### Supplemental figure 10

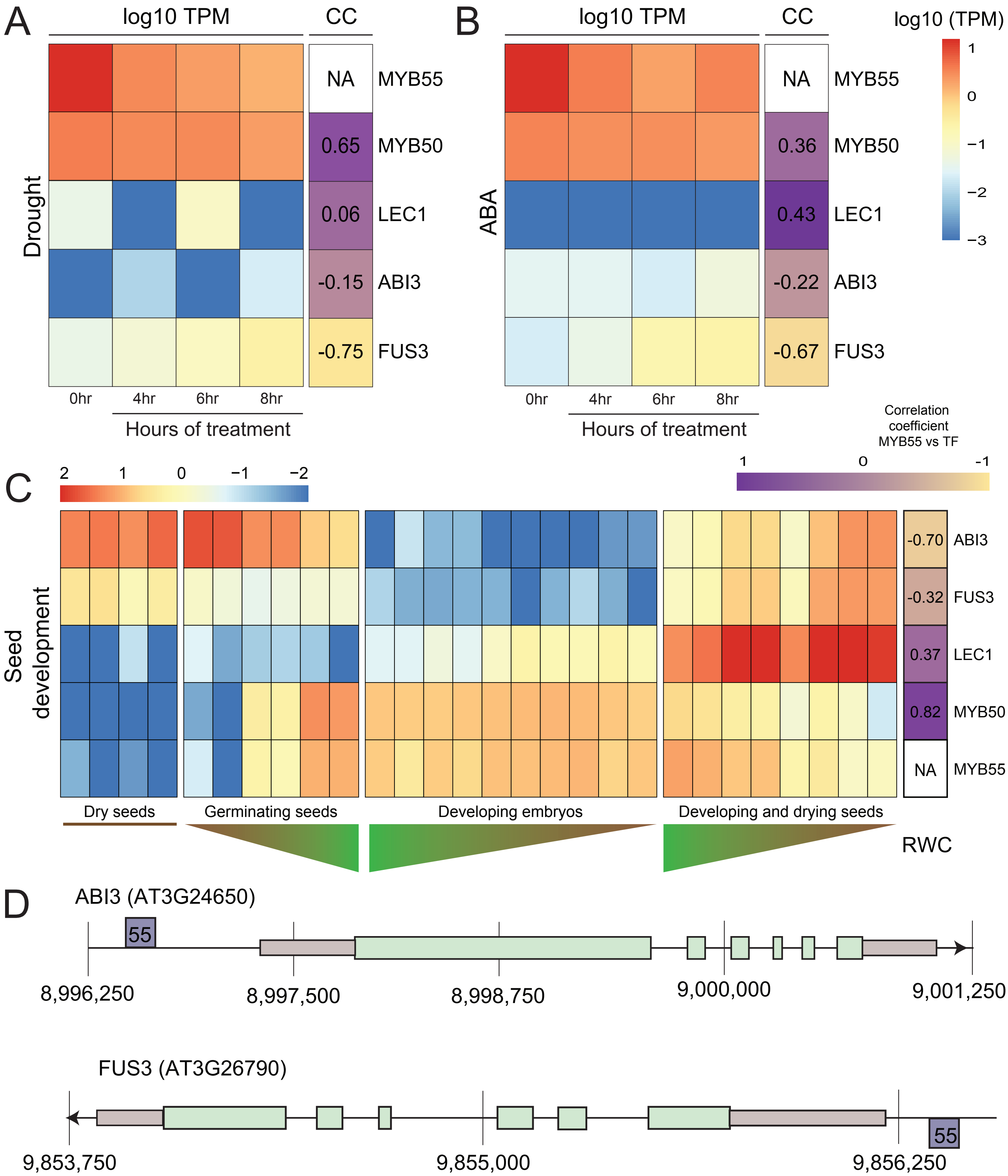

### Supplemental figure 11

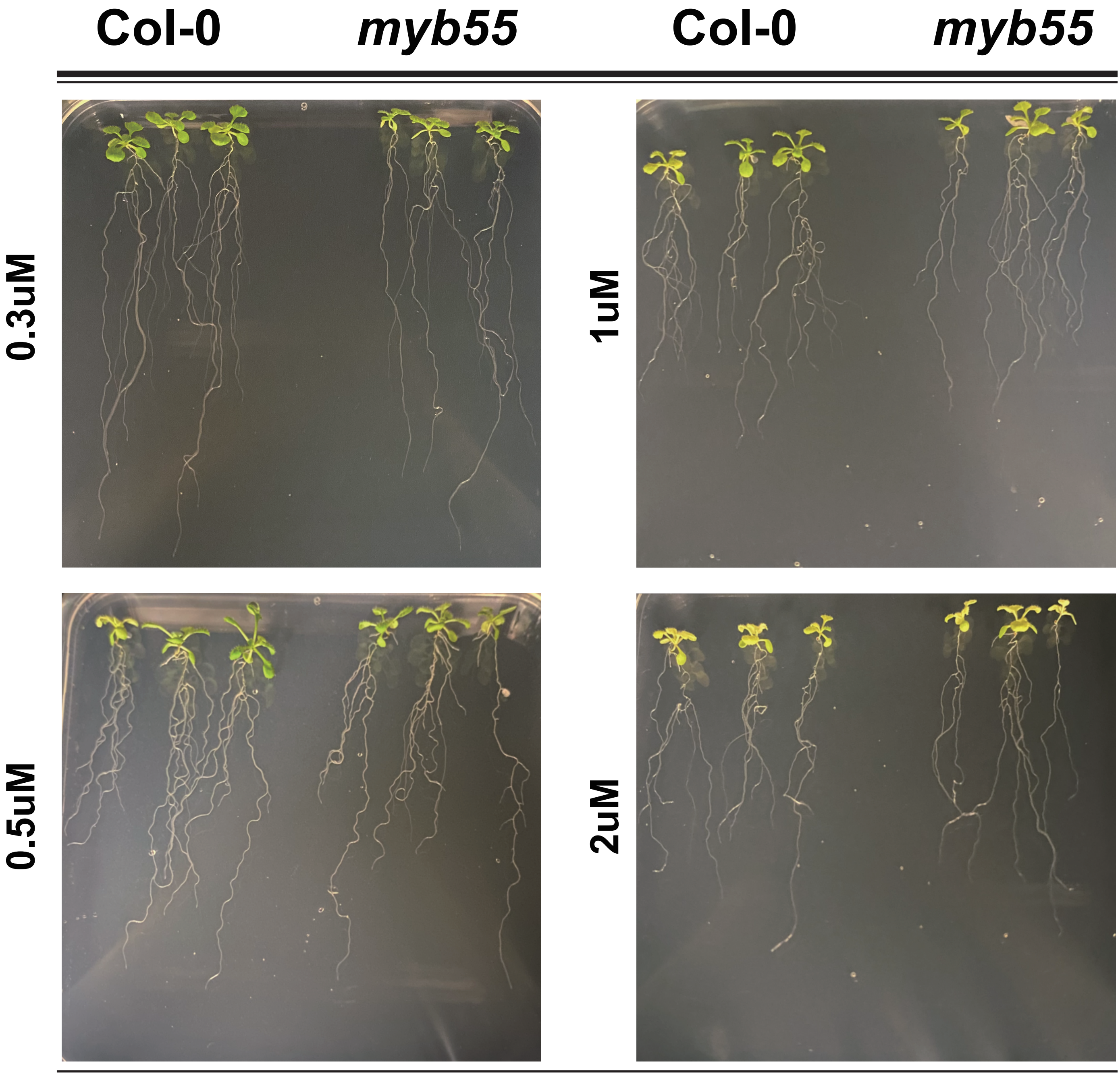
